# Spatially barcoding biochemical reactions using DNA nanostructures unveil a major contact mechanism in proximity labeling

**DOI:** 10.1101/2024.12.17.628942

**Authors:** Zhe Yang, Yu Zhang, Yuxin Fang, Yuan Zhang, Jiasheng Du, Xiaowen Shen, Kecheng Zhang, Peng Zou, Zhixing Chen

**Author notes:** (P.Z.), (Z.C.).

## Abstract

Proximity labeling techniques like TurboID and APEX2 have become pivotal tools in spatial omics research for studying protein interactions. However, the biochemical mechanisms underlying these reactive species-mediated labelings, particularly the spatial patterns of labeling methods within the sub-μm range, remain poorly understood. Here, we employed DNA nanostructure platforms to precisely measure the labeling radii of TurboID and APEX2 through *in vitro* assays. Our DNA nanoruler design enables the deployment of oligonucleotide-barcoded labeling targets with nanometer precision near the enzymes. By quantifying labeling yields using quantitative PCR and mapping them against target distances, we uncovered surprising insights into the labeling mechanisms. Contrary to the prevailing diffusive labeling model, our results demonstrate that TurboID primarily operates through contact-dependent labeling. Similarly, APEX2 shows high labeling efficiency within its direct contact range. In parallel, it exhibits a low-level diffusive labeling towards more distant phenols. These findings reframe our understanding in the mechanism of proximity labeling enzymes, at the same time highlight the potential of DNA nanotechnology in spatially profiling reactive species.

## Introduction

Elucidating protein interaction networks is crucial for understanding fundamental biology and developing new therapeutics. Proximity labeling technology is an emerging yet increasingly popular tool for identifying protein-protein, protein-DNA, and protein-RNA interactions^1-4^. Essentially, proximity labeling involves the site-specific generation of reactive molecules that covalently label biomolecules within a certain distance range, followed by the identification of labeled biomolecules. Since the first conceptual prototypes, BioID^5^ and APEX^6^, the application of various biochemical reactions has led to numerous proximity labeling methods, such as PUP-IT^7^, CAP-seq^8^, μMap^9^, FucoID^10^, PhoXCELL^11^, QMID^12^, CAT-S^13^, BAP-seq^14^, MultiMap^15^, among which APEX2^16^ and TurboID^17^ are the most widely used. These proximity labeling methods provide various options for exploring biomolecular interactions, as their mechanistical difference give rise to varying labeling reactivities and spatial preferences.

Specifically, the radius of proximity labeling is a key parameter determined by the intrinsic reactivity of the active species. Practically, measuring the labeling radius is essential for choosing the right proximity labeling system in various experimental contexts. Despite the widespread use of proximity labeling in protein interactomics, the accurate profiling the distance of proximity labeling has been disproportionally lagging behind due to a lack of molecular tools. There have been a few studies to profile the labeling radius of proximity enzymes^18,19^. For example, the Roux group suggested that the labeling radius of BioID is around 10 nm in cells using a rigid nuclear pore protein complex as a molecular ruler^20^. The MacMillan group developed a method to image the labeling spot of the proximity labeling using super-resolution microscopy^21^ (Figure 1A), concluding that the labeling radius of HRP is 268 nm. While these methods provide valuable spatial information, the *in cellulo* measurements are intrinsically complicated by the cellular environments. Moreover, it is challenging to perform mechanistic studies and establish the full profile of distance-labeling efficiency relationship without *in vitro* reconstructed system of proximity labeling. Without such biochemical parameters on the distance profile, our understanding on the overarching mechanism of proximity labeling remains incomplete.

**Figure 1.**
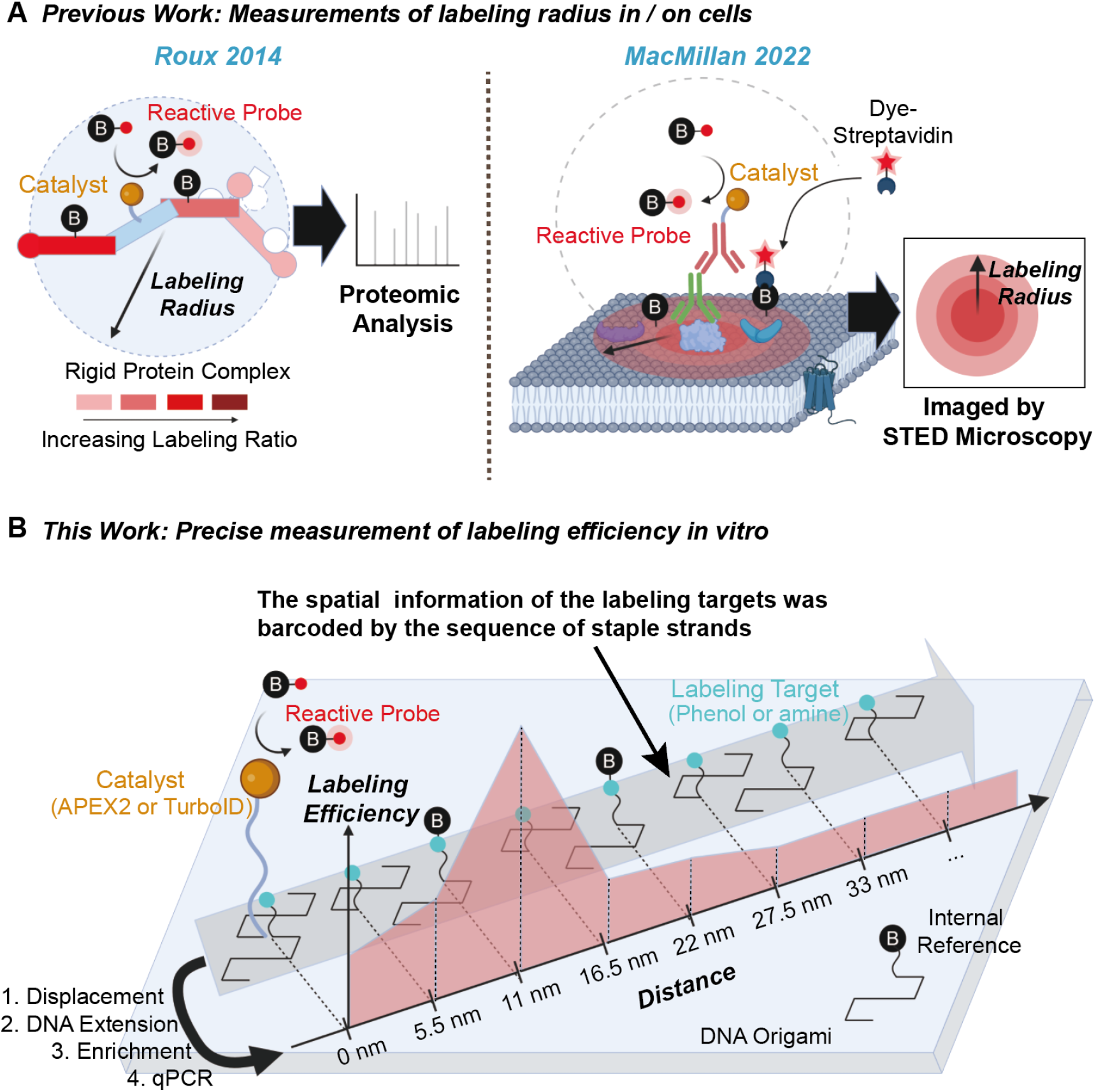
Selected methods for measuring the working distance of proximity labeling. **A)** Labeling radius are previously measured in live cell context using spatial proteomic analysis of a rigid protein complex or super-resolution imaging of membranous targets. **B)** In this work, DNA-nanostructures templated proximity labeling enables accurate profiling of their spatial ranges *in vitro*.

In this study, we devised DNA nanostructures which can accurately template proteins and labeling targets with designated addresses at nanometer accuracy, thereby reconstructing the proximity labeling systems *in vitro*. DNA nanostructures offer the exquisite programmability at the same time ideal distance ranges (1-100 nm) matching the working distances of proximity labeling. Currently, protein-DNA-nanodevices are employed to study various enzymatic cascade reactions by controlling the relative distance and orientation of enzyme components^22-24^. DNA nanostructures can also serve as platforms for single-molecule chemical reactions^25,26^, making them ideal candidates for probing the spatial dynamics of proximity labeling reactions. Our system leverages rigid DNA nanostructures to control the distance between the enzyme catalyst that generates reactive species and the labeling targets, with spatial information of the labeling targets barcoded by the sequence of staple strands. Using pulldown and qPCR quantification to measure labeling efficiency at various distances, we were able to profile the full distance-labeling efficiency map of the APEX2 and TurboID proximity labeling reactions in a controlled microenvironment, which reveals new insights to the mechanism of these proximity labeling reactions.

## Results

### Spatially addressed enzymatic catalyst and labeling targets on DNA origami

We aim to reconstruct the proximity labeling systems *in vitro* to create a chemically-defined environment without unknown factors. Considering the prior knowledge that proximity labeling mainly functions in the nm to μm range, we regard the DNA nanostructures as an ideal molecular ruler for this purpose thanks to their structural rigidity as well as the synthetic programmability. The catalyst, APEX2 or TurboID, can be conjugated to designer DNA oligos with a fusion HaloTag protein. The DNA-Catalyst dyad can be hybridized to a specific address on a DNA origami (Figure 1B). In this setup, the active center of protein catalyst is about 10 nm away from its branching spot on DNA origami. Moreover, selected staple strands were chemically functionalized at the 5’ ends with labeling targets, such as phenols for APEX labeling or amines for TurboID. These functional staple strands were weaved into the DNA origami at designated addresses with known distance to the catalyst. In this way, an array of proximity labeling reactions can be programmed at various distances, whose labeling efficiencies can be then read using standard nucleic acid quantification methods.

To establish a fixed spatial arrangement for APEX2 labeling on DNA origami platform, we first designed modification sites for phenol group at varying distances from the APEX2 site (Figure S1 and Table S2, S3). Then, we installed the phenol groups on the selected staple strands by modifying the amino-modified staple strands with phenol-NHS, as confirmed by mass spectrometry (Figure S2). The phenol-modified staple strands were annealed onto the DNA origami using standard protocols (see methods). Finally, the APEX2-ssDNA strand, prepared separately via the reaction of HaloTag ligand-ssDNA and purified recombinant APEX2-HaloTag fusion protein (Figure S3), were hybridized to the extended handle strand on the DNA origami with a known address (Figure 2A). In this *de novo* designed molecular architecture, both the anchoring site of APEX2 and the phenol group are programmed on the same side of the DNA sheet^27^. Notably, while APEX2 can label guanosine bases, the phenoxyl free radical shows substantially lower reactivity toward double-stranded DNA compared to its preferred substrate, phenol^28,29^.

**Figure 2.**
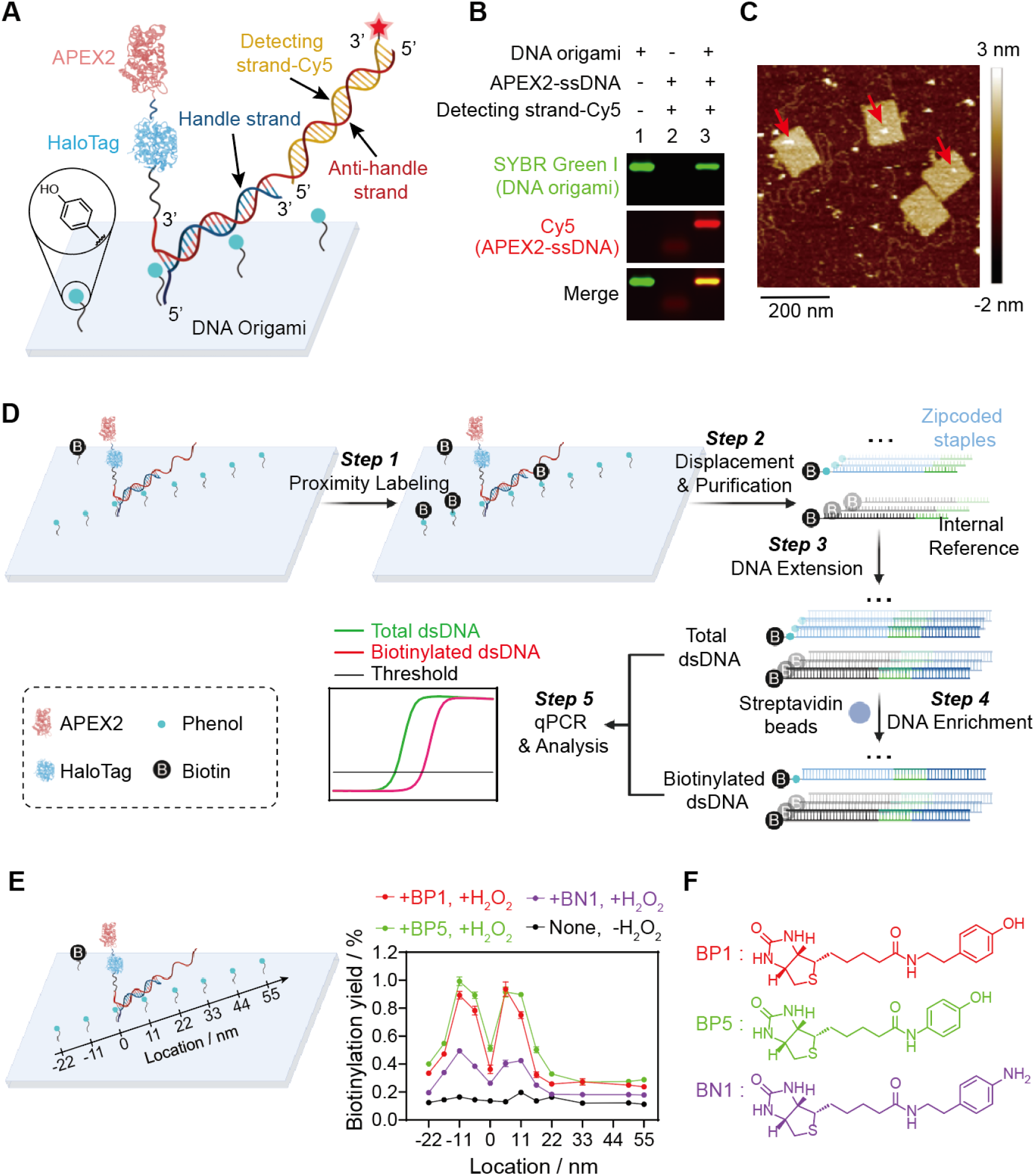
Measuring the labeling radius of APEX2 with DNA origami. **A)** Schematic diagram of assembly of DNA origami and APEX2. **B)** Gel characterization of the assemble of DNA origami and APEX2. **C)** AFM image of DNA origami-APEX2 assembles. **D)** Workflow of labeling efficiency measurements at destinated addresses. After performing proximity labeling reaction, the DNA oligos are purified, elongated, enriched, and subjected to qPCR quantifications. Benchmarking with an internal reference gives the labeling yields at different locations. **E)** Plot of biotinylation yield against phenol location. **F)** Chemical structures of biotin-phenol/aniline substrates.

To confirm the assembly of the APEX2-origami complex, a detecting strand modified with Cy5 at the 3’ end was hybridized to the anti-handle strand covalently conjugated to the APEX2 fusion protein. Gel electrophoresis followed by in-gel fluorescence imaging indicated a successful anchoring of APEX2 to the origami (Figure 2B). AFM images further confirmed that APEX2 was precisely positioned to the designated address on the origami (Figure 2C and Figure S4).

Following the assembly of APEX2 onto the DNA origami, proximity labeling was initiated by the addition of biotin-phenol and hydrogen peroxide (Figure 2D). After 30 min, the reaction was quenched by the addition of trolox (10 mM) and sodium ascorbate (10 mM). The staple strands and scaffold strand in the origami were then peeled off by a strand-replacement reaction using the F-5.2 primer and Vent polymerase (Figure 2D and Figure S5A), followed by a purification step (Figure S5B, D. see methods for detail). Since the staple strands have similar base lengths (32 bp) to microRNAs, we used a protocol typically employed for detecting miRNA to determine the biotinylation ratio of the modified staple strands^30^. The purified staple strands with poly(T)_25_ were extended to an appropriate length (90 bp) using an adapter strand containing poly-A (Figure S5C, F). Subsequently, the extended staple strand, referred to as Input, was enriched using streptavidin-beads^8^, resulting in an enriched sample referred to as Enrich (Figure 2D). Notably, we introduced a constitutively biotinylated staple strand as an internal reference in the very beginning to control the loss during the enrichment process.

Moving to the analysis step, these samples were subjected to qPCR using specific primers that do not interfere with each other (Figure S6). By comparing the Ct values of Input and Enrich, we calculated the biotinylation efficiency (F) of each phenol group-modified staple strand using **Equation 1**, where E_staple_ is the amplification efficiency of each staple (Figure S7 and Table S1), Ct_staple_ Input and Ct_staple_ Enrich are Ct value before and after enrichment, respectively. Similarly, E_IR_, Ct_IR_ Input, and Ct_IR_ Enrich are the amplification efficiency of internal reference, Ct value before and after enrich, respectively.

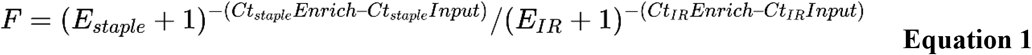

With the origami template and the DNA analysis protocol, the labeling efficiencies of phenols were plotted against their distance to APEX2 (Figure 2E, F and Figure S8). The results indicate that significant labeling occurs within 55 nm, with the site within ±11 nm showing the highest labeling. Screening of alternative substrates with different reactivities gave different labeling efficiencies yet similar distance profiles^29,31^. The control group without substrate and hydrogen peroxide give a low yet consistent signal (<0.2%), indicating a relatively clean pull-down background. Notably, inter-molecular labeling efficiency is also measured by the co-addition of an empty DNA origami with orthogonally barcoded phenols but without a catalyst (Figure S9A, B). At the origami concentration of 10 nM, the labeling at adjacent origami were very low, comparable to the pull-down background. To simulate the intracellular labeling of APEX2, cysteine was added to the buffer to create a reducing environment. Interestingly, the labeling efficiency of APEX2 is generally decreased, but the overall trend over various distance remained the same (Figure S10). The labeling efficiency of APEX2 in 98% of heavy water showed a similar drop.

Overall, APEX labeling at DNA origami template gave two features. 1. Expectedly, it is highly distance-dependent, with intermolecular samples essentially unlabeled and strong labeling within 70 nm. 2. Paradoxically, labeling around 10 nm gave an abruptly high signal, which is even stronger than that at the point of origin.

### dsDNA is a fine ruler for measuring the distance of APEX2 labeling

We were intrigued by the APEX2 labeling behavior in the short range within 20 nm, where the maximal labeling happened at around 10 nm on origami. As it has become impractical to measure on origami with smaller step size, we turned to a double-stranded DNA (dsDNA) scaffold system to fine tune the distance between the enzyme and the labeling targets with base pair-level precision. Briefly, a phenol group was labeled at the 5’ of a single-stranded DNA, while APEX2 was labeled at the 5’ or a designated position in the middle of its complementary strand (Figure 3A, B and Figure S11A, B). Due to the rigidity of dsDNA with a persistence length of 50 nm^32,33^, when the two complementary DNAs hybridized to form a double strand, APEX2 and the phenol group can be well separated at designated distances. In this setup, the active center of APEX2 in the dsDNA system was about 7 nm away from its conjugation site on DNA. By adjusting the anchoring position of the APEX2 on the dsDNA structure, the distance to the phenol group could be modulated with 1 nm step size (Figure 3A).

**Figure 3.**
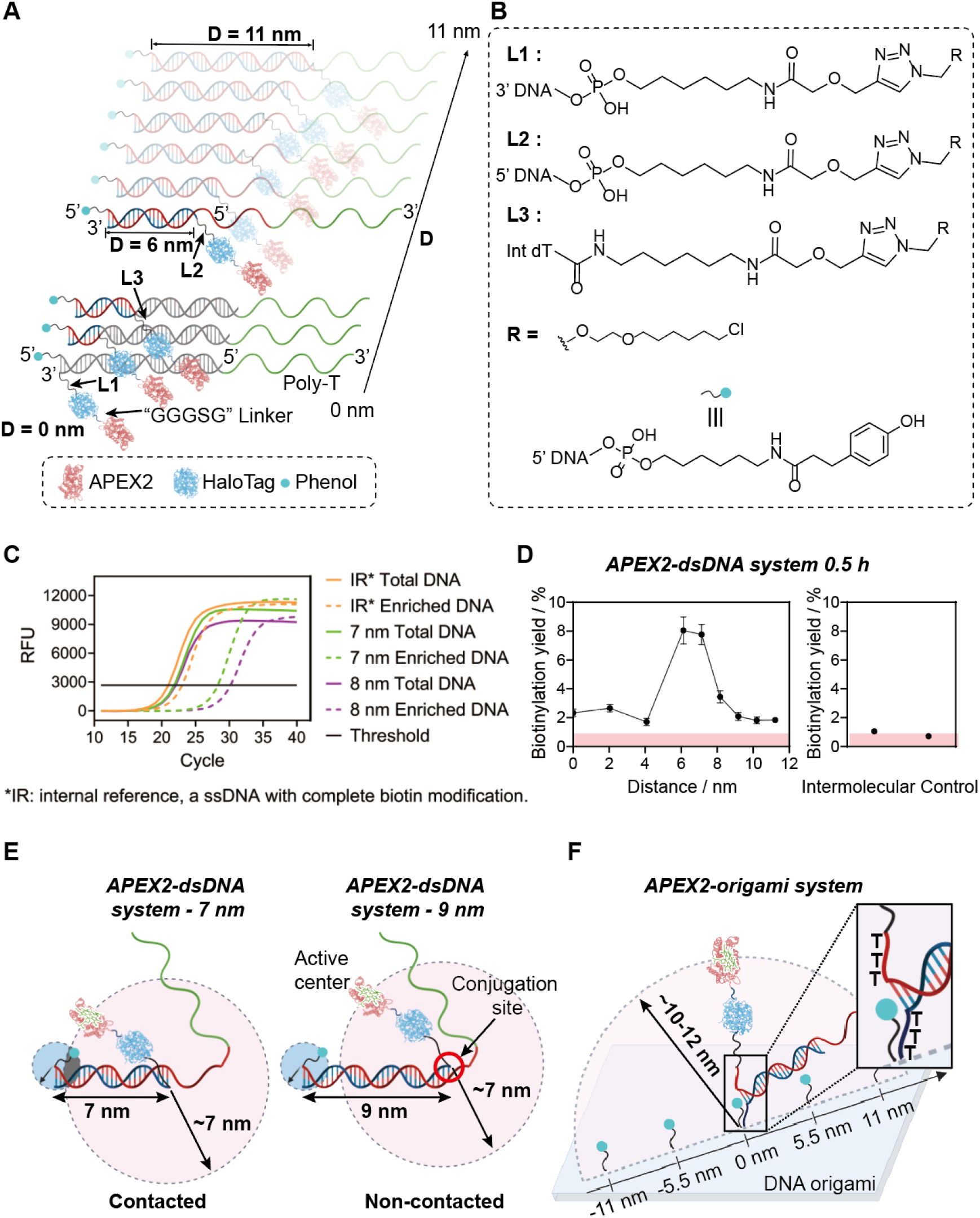
dsDNAs as finer nanorulers to accurately profile the labeling radius of APEX2. **A)** Schematic diagram of dsDNA-templated APEX2 with phenol groups at different distances. D refers to the distance between phenol and the conjugation sites of APEX2 on dsDNA. **B)** Chemical structures of linkers in **A. C)** Representative qPCR curves for quantifying labeling efficiencies. **D)** Labeling efficiencies of APEX2 to phenol groups with different distances spaced by dsDNA. Pink areas represent the level of background intermolecular labeling. **E)** Schematic diagram of the contact-dependent labeling model (left) and the non-contact-dependent labeling model (right) of APEX2 in dsDNA system. **F)** Schematic diagram of APEX2 on DNA origami, showcasing a longer contact labeling radius attributed to longer triple T linkers.

Using this setup, the APEX2-dsDNA-phenol group systems with varying distances from 0 to 11 nm (Table S6, S7) were pooled in the same tube. The labeling reaction was initiated by adding biotin-phenol and hydrogen peroxide. Following a 30-min incubation, the subsequent steps including double-strand extension, enrichment, and qPCR were performed to calculate the labeling efficiency of APEX2 on the phenol group across different distances (Figure S12). The APEX2 - dsDNA system gave consistent results compared to that of the APEX2 - DNA origami system, demonstrating high labeling efficiency within 12 nm (Figure 3C, D). In this setup, the highest labeling efficiency (8%) took place at 6 nm and 7 nm, which was approximately fourfold greater than that at other positions (2%). In intermolecular control, the labeling yield was the lowest (1%). Notably, there appeared to be sharp drops between the peak at 6-7 nm and nearby spots at 4 or 8 nm. The dsDNA results corroborate the origami results, suggesting a labeling mechanism that deviates from simple diffusion of reactive phenol radicals, which should give a monotonic decay in labeling yield over distance.

### APEX labeling has a proximity labeling portion and a contact labeling portion in parallel

It has been suggested that APEX2 labeling occurs through the interaction of the released phenoxyl radicals with surrounding phenol groups. Supposedly, the labeling efficiency should be determined by the local concentration of phenoxyl radicals diffused from the active center of APEX2. We use Brownian motion to simulate the distance-dependent diffusion of phenoxyl radicals generated at the APEX2 active center. The concentration change of phenoxyl radicals is described by **Equation 2**, where *n*(*r,t*) is the concentration of phenoxyl radicals at a distance *r* from the APEX2 active center, *D* is the diffusion coefficient, and *t* is the diffusion time^34^.

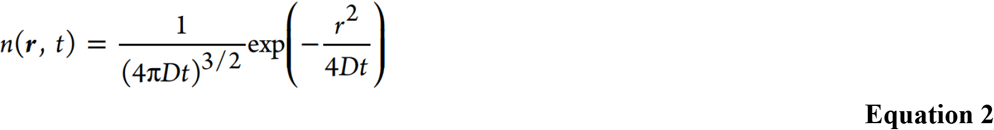

A prior study has suggested a half-life of 0.1 ms for phenoxyl radicals in the aqueous environment and in the absence of quenchers such as glutathione^35^. Starting with a reported diffusion coefficient of phenol at 1000 *μ*^2^/*s*^36^, we inferred the diffusion coefficient of the biotin-phenoxyl radical to be about 200 *μ*^2^/*s* based on their molecular weights^37^. Our simulation results reveal that the diffusion radius is within 500 nm (Figure S13A), aligns with labeling radius measured with STED imaging which is around 268 nm^21^. Our simulation also indicated that the concentration of phenoxyl radicals decreases slightly (less than 6%) within 70 nm, suggesting that the labeling efficiency of APEX2 remain relatively constant within short distance, which aligns with our findings within 22-66 nm of the origami system and 9-11 nm in the dsDNA system (Figure 2E, Figure 3D and Figure S9). However, the high labeling efficiency observed at 5.5-11 nm in the origami system and 6-7 nm in the dsDNA system contradicts the diffusion model, indicating an alternative labeling mechanism beyond phenoxyl radical diffusion labeling.

Taking the size of APEX2 (Figure S14)^38^ and the linker length into account, the distance between the active center of APEX2 and the DNA conjugation site is approximately 10-12 nm in the APEX2-DNA origami system. Consequently, the phenol group positioned at 5.5 nm and 11 nm away in the APEX2-DNA origami system can reach the APEX2 active center with a preferred conformation (Figure 3F). Similarly, the phenol group at 6 nm and 7 nm in the APEX2-dsDNA system are also capable of direct contact labeling with the APEX2 active center (Figure 3E). Therefore, we attribute the enhanced labeling efficiency at 5.5-11 nm in the origami system and 6-7 nm in the dsDNA system to the direct oxidation of phenol with the Fe^IV^ heme complex at the active center of APEX2, which is significantly faster than a relay oxidation from a heme-sensitized phenoxy radical. The resulting phenoxyl radical would then engages in diradical recombination with the biotin-phenoxyl radicals present in the solution^39^, finishing the biotin labeling. Notably, while the APEX2-HaloTag protein remains the same, in the dsDNA system, the redundant triple T linkers in the origami system were removed (Figure 3E), and the chemical linkers for click chemistry was further optimized (Figure S3 and Figure S11), giving a shorter effective linker than that in the origami. Such tuning in linkers matched exactly with the difference in labeling radius (6 - 7 nm on dsDNA vs 5 - 11 nm on origami), further supporting the contact mechanism. In summary, our findings inferred that APEX2 exhibits a dual labeling mechanism, featuring both a previously appreciated diffusive labeling process that covers the distal phenols within 500 nm, and a newly unveiled strong contact labeling within 10 nm.

### Measuring the distance of BioID and TurboID using dsDNA ruler

We further employed DNA nanoruler to investigate the labeling distance and mechanism of BioID and TurboID, both of which are engineered enzymes derived from the biotin ligase BirA. Previous studies have shown that BioID can label proteins within a distance of 10 nm in cells^5,20^. Therefore, we chose the fine ruler of dsDNA system functioning within 11 nm to measure the labeling distance of these two enzymes.

TurboID can modify lysine residues of proteins and aliphatic primary amines on oligonucleotide^40^. Therefore, we adapted the dsDNA scaffold from the previous setup, replacing phenol groups with amino groups to mimic lysine residues, and substituted APEX2 with BioID or TurboID (Figure S15). The conjugation of BioID and TurboID to single-stranded DNA (ssDNA) was confirmed via SDS-PAGE (Figure S16). The reaction buffer was changed to sodium tetraborate buffer (0.1M Na_2_B_4_O_7_, pH 8.5) to deprotonate and enhance the reactivity of the amino groups. Proximity labeling was initiated by the addition of biotin and ATP. After the labeling reaction, Tris (10 mM) was added to terminate the reaction, and the labeling efficiency was assessed using qPCR following the same workflow (Figure S12).

In the BioID-dsDNA system, there is a highest labeling efficiency of ∼0.1 % at 6 nm after an 18-hour reaction (Figure 4A, B), with undetectable labeling at shorter times (Figure S17). The pull-down background, which indicated the non-specific absorption of unlabeled DNA by streptavidin beads, are only ∼0.028(±0.001)%. The intermolecular labeling of BioID to another amino-ssDNA (2 nM in solution) is even less (∼0.010(±0.004)%). The labeling efficiency at 6 nm is significantly higher than the pull-down background while the labeling at other distances are comparable to intermolecular labeling. In the TurboID-dsDNA system, we observed a high labeling efficiency within 6 nm after a reaction time of 0.5 hours. However, the labeling efficiency dropped sharply beyond 6 nm, approaching background levels (Figure 4A, C). Similar results were observed for extended labeling times (Figure S18). These results are consistent with the previous studies that TurboID has a higher labeling efficiency than BioID^17^. Interestingly, the labeling efficiency dropped significantly at 7 nm in both the BioID-dsDNA system and the TurboID-dsDNA system, which deviates from the previous perceived labeling mechanism. Moreover, labeling within 6 nm are paradoxically less efficient than that at 6 nm. We speculate that the labeling of BioID and TurboID function through a contact-dependent mechanism instead of the diffusion of active biotin-5’-AMP: at 6 nm, BioID and TurboID holding the biotin-5’-AMP can physically reach the amino group with an optimal geometry (Figure S19A), resulting in a highest labeling efficiency. Labeling within 6 nm is sterically unfavored, while labeling beyond 6 nm are out of range, resulting in negligible labeling (Figure S19B).

**Figure 4.**
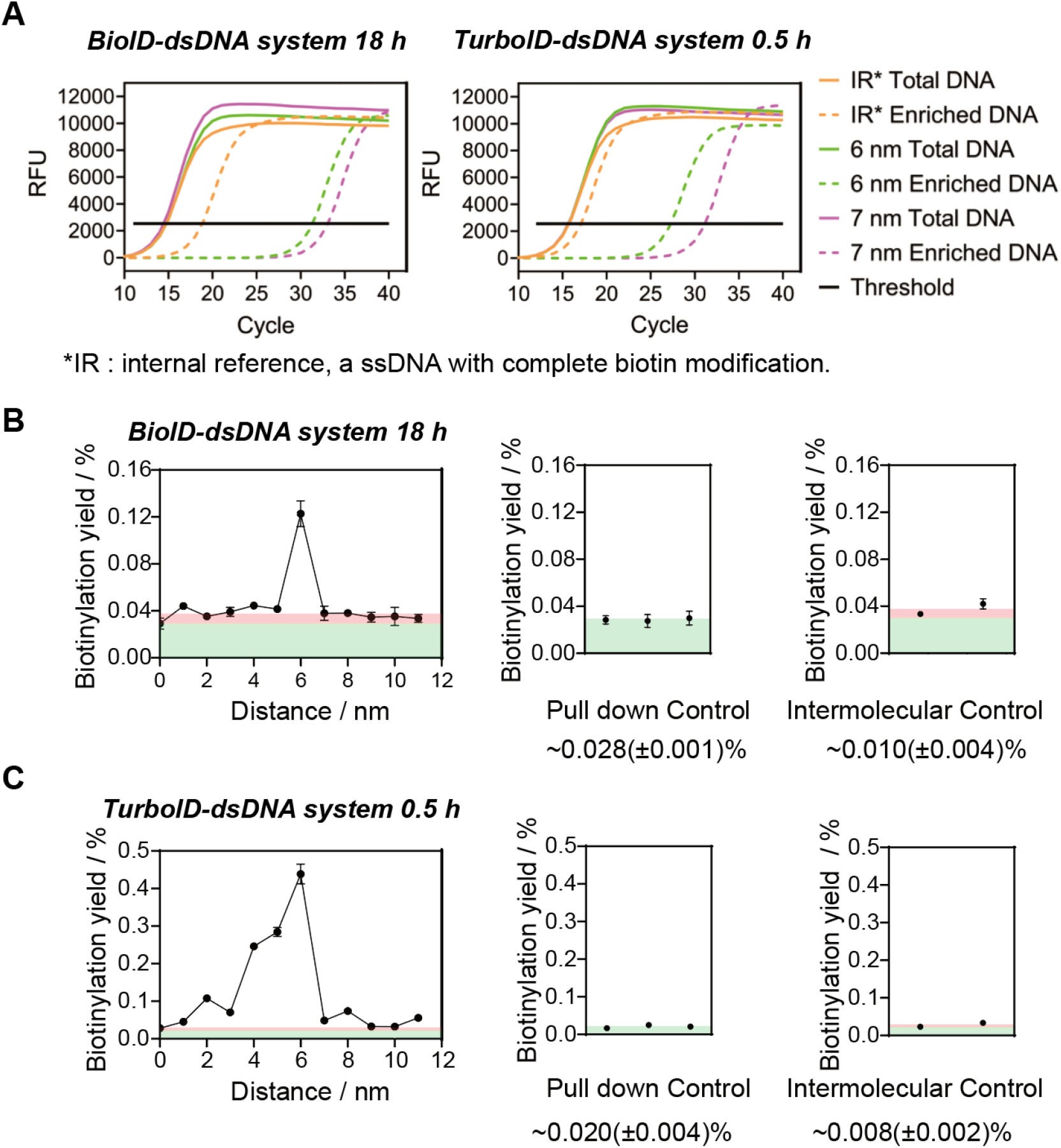
Measuring the labeling radius of BioID and TurboID by dsDNA-nanorulers. **A)** Representative qPCR curves for quantifying labeling efficiencies of BioID and TurboID to amines with different distances spaced by dsDNA. **B)** Biotinylation yield-distance plot of BioID. **C)** Biotinylation yield-distance plot of TurboID. Pink areas represent intermolecular labeling background and green areas represent pull down background.

### Enzymology of TurboID supports a contact-labeling mechanism

It is previously suggested that biotin-5’-AMP is first generated by TurboID using ATP and biotin as precursors, followed by the release of biotin-5’-AMP which labels distal amines. Our data from the DNA ruler, however, suggested a strong preference on contact labeling. To further understand the labeling profile of TurboID, we measured its affinity with biotin-5’-AMP. Following an established protocol^41^, we chemically synthesized biotin-5’-AMP and determined its association and dissociation rate constants with BioID and TurboID using stopped-flow spectroscopy (Figure 5A, B, C and Figure S20). The association between TurboID and biotin-5’-AMP is as fast as that of BirA and BioID (Figure 5D). Surprisingly, the dissociation kinetic (*k*_*off*_) of BioID and TurboID are relatively close to each other. Consequently, the dissociation constant *K*_*D*_ for the BioID·biotin-5’-AMP complex was measured to be 1.6 × 10^−8^ M (Figure 5D, Figure S20 and Figure S21), while TurboID·biotin-5’-AMP exhibited a *K*_*D*_ of 4.0 × 10^−8^ M (Figure 5D). The measured *K*_*D*_ of TurboID · biotin-5’-AMP and BioID · biotin-5’-AMP is comparable to that reported for BirA R118G in a previous study^41^. Upon generation of biotin-5’-AMP in the pocket of BioID and TurboID, the lifetime of the complex is at the order of 10 s before the dissociation.

**Figure 5.**
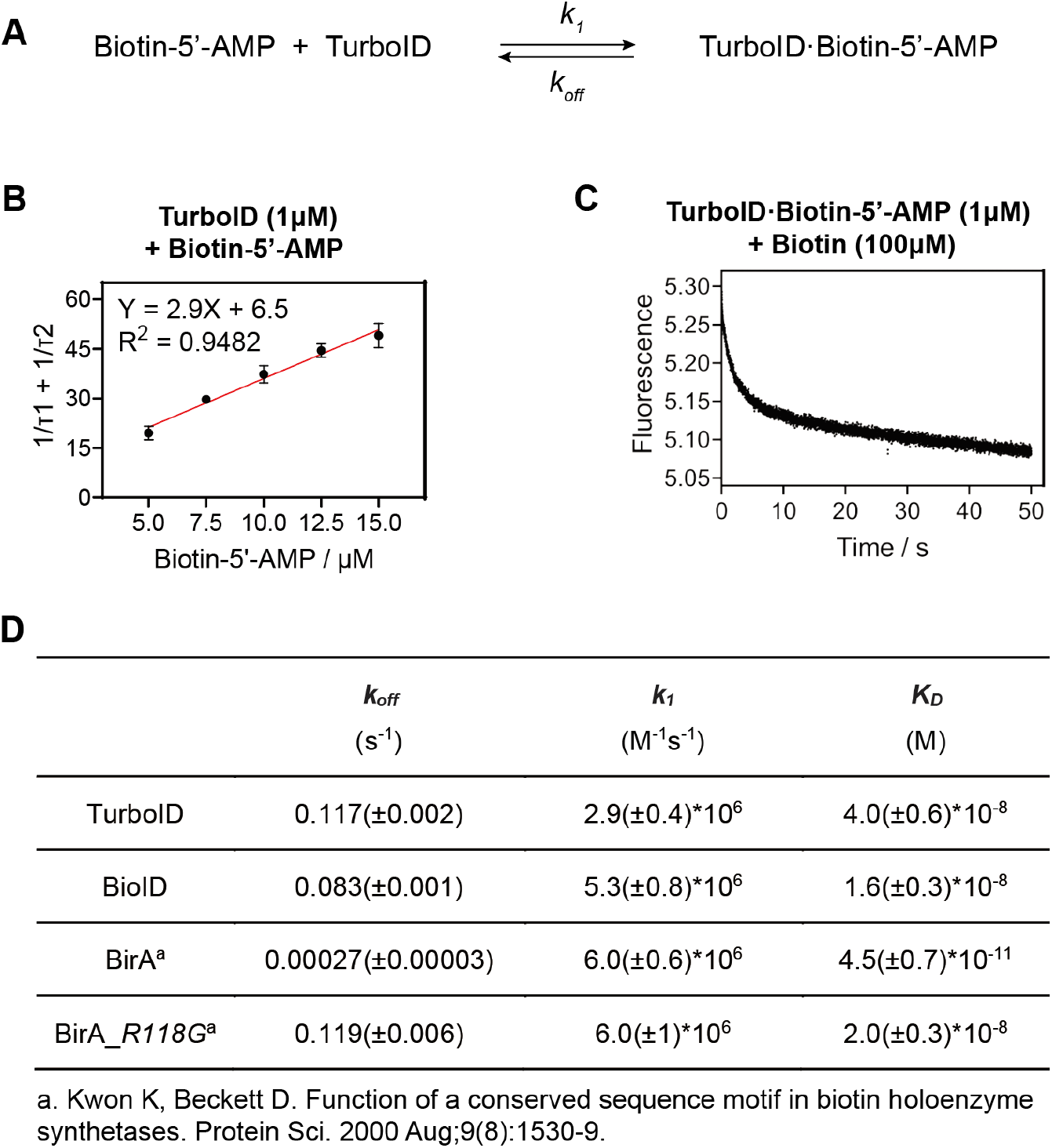
Enzymatic parameters of TurboID. **A)** Chemical equation of association and dissociation of TurboID with biotin-5’-AMP. **B)** Dependence of the sum of the apparent rates (1/τ1 and 1/τ2) of the two kinetic phases on ligand concentration for TurboID, where τ1 and τ2 are the relaxation times for the first and second kinetic phases^41^, respectively (see methods). **C)** Dissociation of the complexes of TurboID bound to biotin-5’-AMP measured by stopped-flow fluorescence. **D)** Kinetic and thermodynamic parameters of biotin-5’-AMP binding to TurboID, BirA, and BioID.

As TurboID was devised with drastically higher labeling efficiency than BioID, the underlying mechanism, however, was elusive. One speculation from the original TurboID paper was that it offers faster dissociation with biotin-5’-AMP^17^. However, our data, showcasing TurboID is only marginally prone to biotin-5’-AMP release than BioID, is against this speculation. Coming to the distance profile, BioID and TurboID gave consistent labeling preference of reachable amines, suggesting that most of the labeling reactions between biotin-5’-AMP and amines happened within the BioID/TurboID pocket thanks to the relatively long dissociation lifetime (∼ 10 s). Overall, the enzymology of TurboID revealed a tight binding with biotin-5’-AMP, supporting the contact-labeling mechanism unveiled with DNA rulers.

## Conclusion and Discussion

In this study, we developed a DNA nanostructure-based platform to precisely control the distance between enzymes and labeling targets, allowing the reconstruction of proximity labeling systems *in vitro*. By employing qPCR to measure the relative labeling efficiency at different distances, we elucidated new aspects of the proximity labeling mechanism that challenges the current paradigm. Our results suggest that APEX2 labeling involves both contact and non-contact mechanisms (Figure 6A), while BioID/TurboID labeling is primarily through a contact-based mechanism within the reach of ligase (Figure 6B). Such universal contact-based labeling is substantiated with solid evidence using DNA rulers. From a proteomics perspective, our findings provide practical guide into selecting appropriate proximity labeling methods as well as insights for data interpretation. TurboID is recommended for more accurate identification of interacting proteins due to its strict contact-dependent labeling, while APEX2 is better suited for longer-distance labeling.

**Figure 6.**
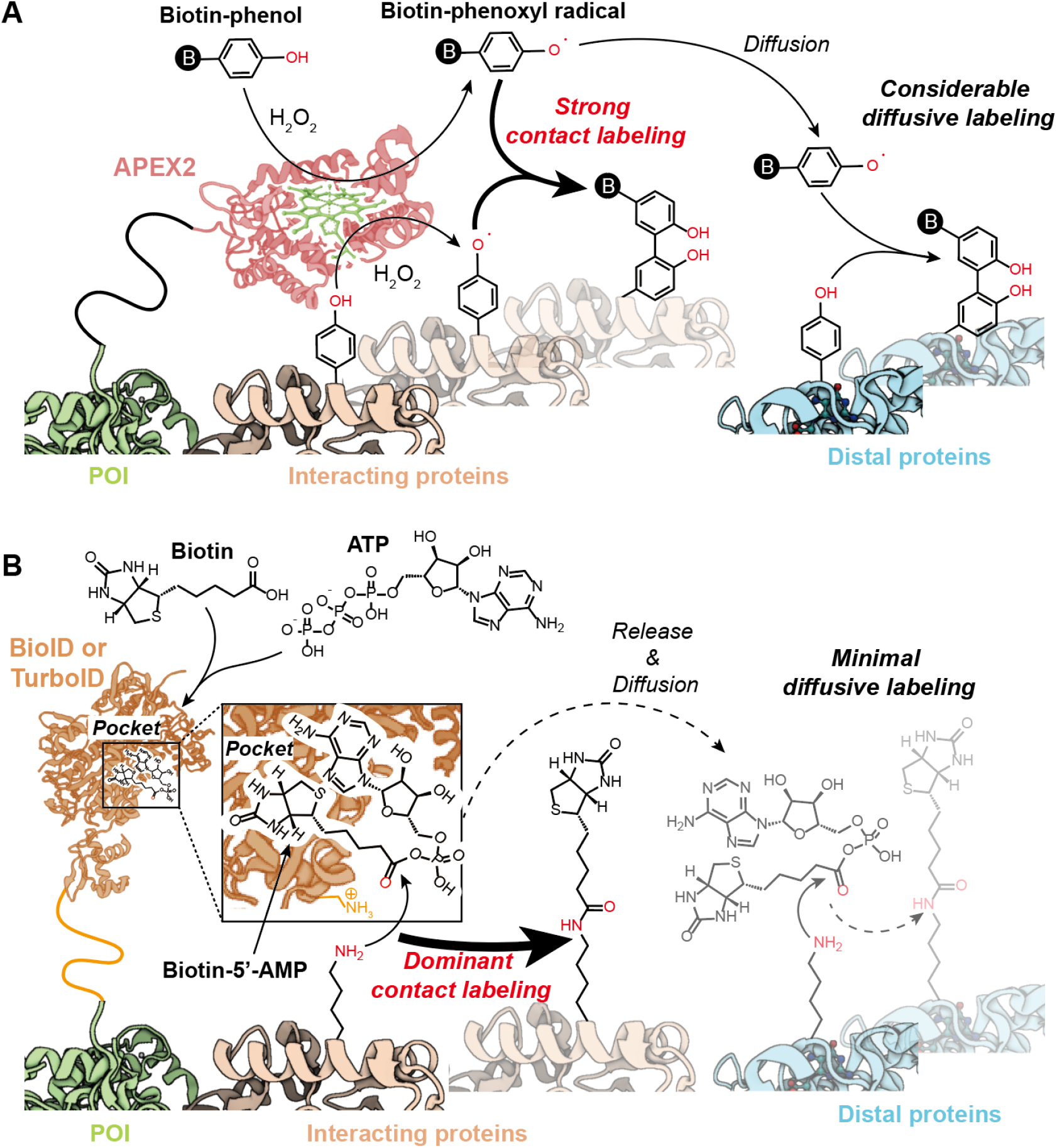
Proposed labeling profile of APEX2 and TurboID, whose distinctive mechanisms give rise to different spatial preference. **A)** APEX2 involves a combination of contact and diffusive labeling. In parallel to the widely appreciated phenol radical-mediated diffusive labeling, the proximity-induced oxidation of adjacent phenols directly from the heme is favored, giving a strong contact labeling which has been largely neglected in previous mechanistic researches. **B)** BioID and TurboID function primarily through contact-dependent labeling, where the generated biotin-5’-AMP was mainly consumed by adjacent amino groups from interacting proteins before released from the pocket, giving little diffusive labeling at distal proteins. POI: protein of interest.

Mechanistically, this work addresses several long-standing controversies: First, the half-life of biotin-5’-AMP at neutral pH should be minutes^42^, it would have rendered BioID/TurboID a long-range proximity labeling method if they function through diffusive labeling mechanism (Figure S22). However, multiple works have suggested that BioID and TurboID function within 10 nm^1,20^. The working model proposed in this work, where BioID/TurboID functions through the direct labeling of adjacent amines with BioID/TurboID·biotin-5’-AMP complex, reconcile with the previous data showing that BioID/TurboID labeling happens predominantly within 10 nm. Second, for APEX/peroxidase labeling, while electron microscopy suggested ∼20 nm labeling resolution^6,18^ yet STED microscopy gave a labeling radius of ∼270 nm^21^, our data provide a unified model. The heavy signal within 20 nm can be attributed to the direct contact sensitization from heme, where the diffusive labeling of phenol radical gives a small portion of distal signal with a radius of ∼270 nm. Overall, this work adds to the few approaches to measuring the distance parameter of proximity labeling. The *in vitro* labeling patterns herein unveil previously uncharted mechanistic insights which reconcile the existing evidences obtained using different approaches.

From the view of chemical tools, the nanoruler platform we established through DNA nanostructures proves versatile for measuring the diffusion of active molecules at the nanoscale. Complementing the FRET dynamic ranges of 1-10 nm^43^ and super-resolution imaging which offer precise measurements at >20 nm^44,45^, this piece exemplified the power of DNA rulers for profiling microscopical patterns in the 1-100 nm window^46,47^. Notably, this work integrates a qPCR-based DNA quantification scheme with the DNA origami platform, capitalizing on the precise addressability of DNA origami structures. This work opens new avenues for investigating biochemical reactions within microenvironments and provides fresh insights into detecting complex reaction intermediates. While earlier studies rely on atomic force microscopy (AFM) to image single-molecule reactions on DNA origami platforms^25,26^, we advocate to introduce detection methods from modern molecular biology such as qPCR with boosted sensitivity. We foresee a broad potential for DNA nanotechnology to serve as a robust quantitative tool in biochemical research.

## Acknowledgments

This study was supported by Li Ge-Zhao Ning Science Youth Research Foundation, the Center for Life Sciences (CLS) of Peking University, the Ministry of Science and Technology (2022YFA1304700), the National Natural Science Foundation of China (32088101), and Beijing National Laboratory for Molecular Sciences (BNLMS-CXXM-202403). PZ is sponsored by Bayer Investigator Award. We thank National Center for Protein Sciences at Peking University in Beijing, China, particularly Dr. Siying Qin, for technical assistance with AFM. We thank Profs. Alice Ting, Profs. Boyuan Wang, Dr. Jing Ling, Dr. Tianyan Liu, Dr. Yi Li, and Dr. Xinyue Zhou for helpful discussions.

## Author Contributions

Zhixing Chen, Peng Zou, and Zhe Yang designed research. Zhe Yang preformed the research. Yu Zhang, Yuxin Fang, Xiaowen Shen and Kecheng Zhang analyzed part of the data. Yuan Zhang and Jiasheng Du synthesized the biotin-5’-AMP. Zhe Yang and Zhixing Chen wrote the manuscript with input from all authors.

## Competing Financial Interests

We declare that none of the authors have competing financial interests.

## Notes

### Competing Interest Statement

The authors have declared no competing interest.

