## Supplementary information for "Spatially barcoding biochemical reactions using DNA nanostructures unveil a major contact mechanism in proximity labeling"

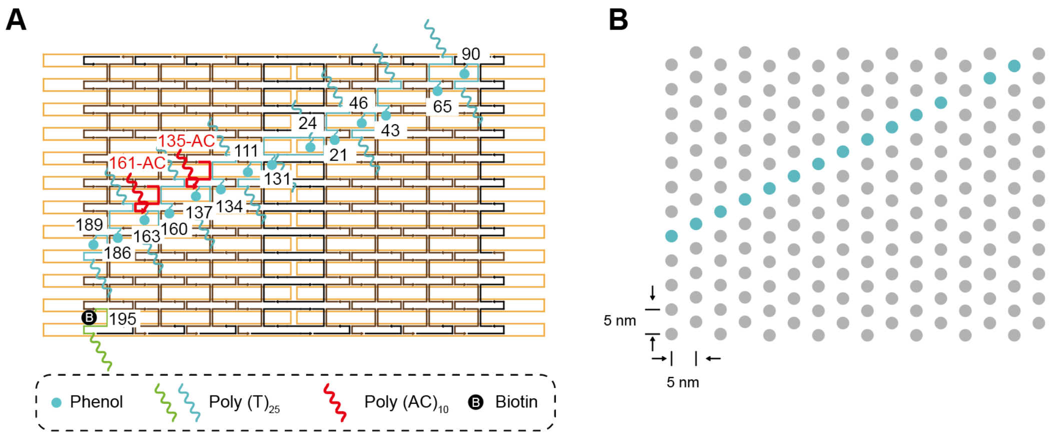


**Figure S1. Design of phenol group sites, APEX2 anchoring sites, and internal reference site on DNA origami. A)** Design details of the phenol group modification sites (blue), APEX2 anchoring sites (red), and internal reference site (green) on DNA origami. The blue dots refer to phenol. The blue and green wavy lines represent poly $(T)_{25}$, and the red wavy lines represent poly $(AC)_{10}$. The sequences of specific staple strands are shown in Table S2, S3, following their numbers. **B)** A simplified design diagram of the DNA origami in A.

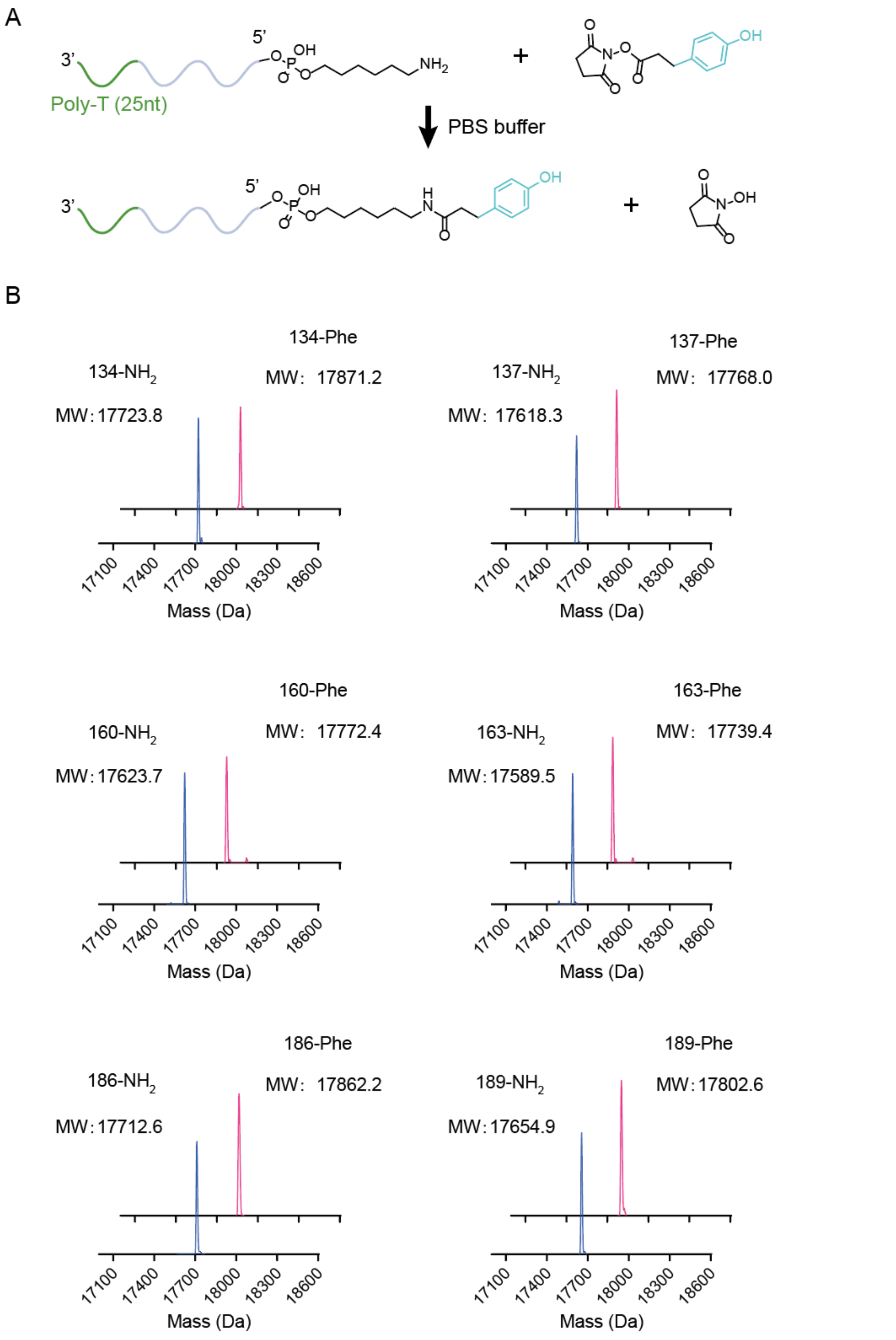


**Figure S2. Characterization of ssDNA- phenol group conjugate. A)** Schematic diagram of ssDNA conjugated with phenol group. **B)** Mass spectrum of ssDNA - phenol group conjugate.

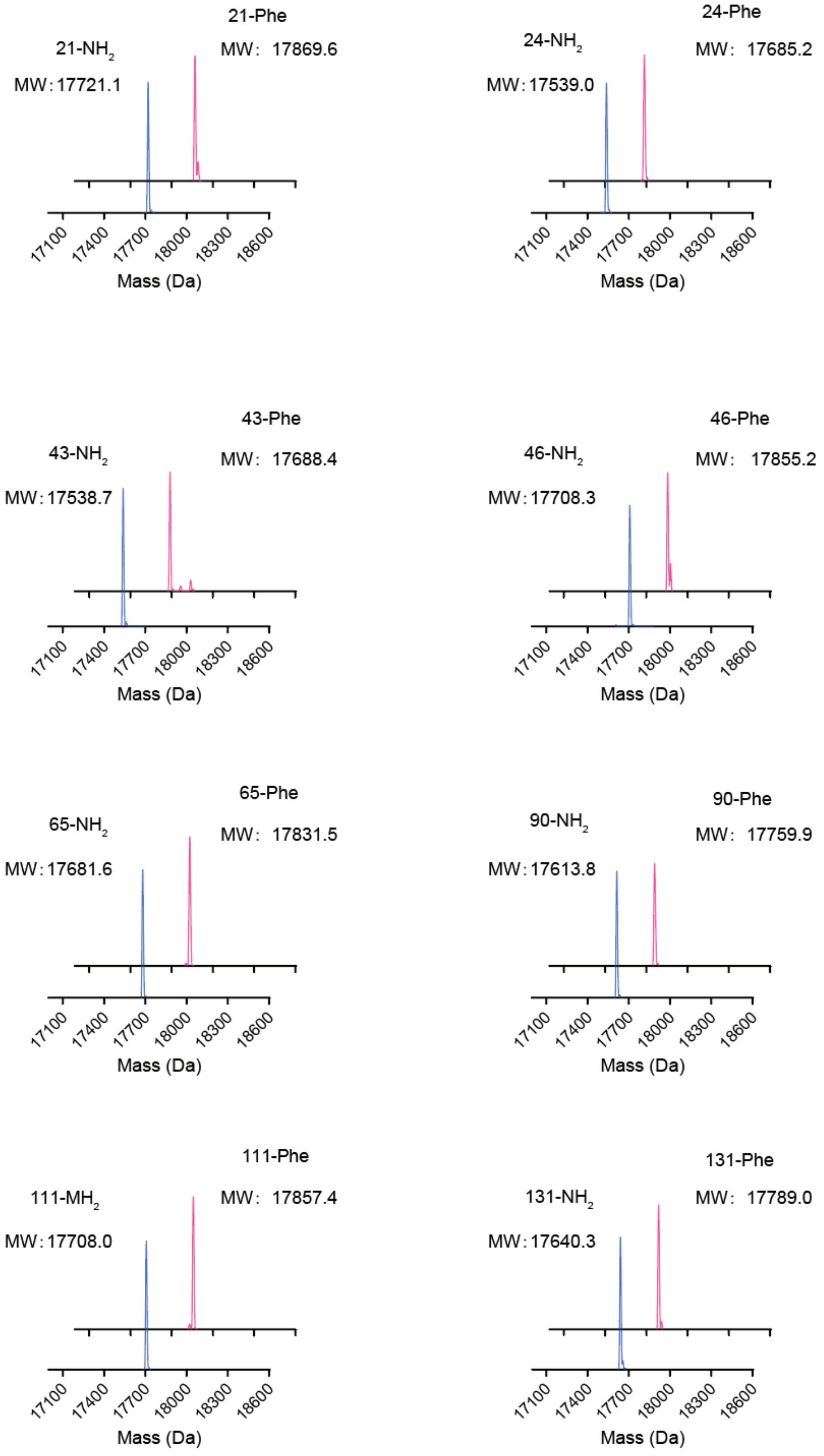


**Figure S2. Characterization of ssDNA-tyrosine residue conjugate. A)** Schematic diagram of ssDNA conjugated with tyrosine residue. **B)** Mass spectrum of ssDNA - tyrosine residue conjugate.

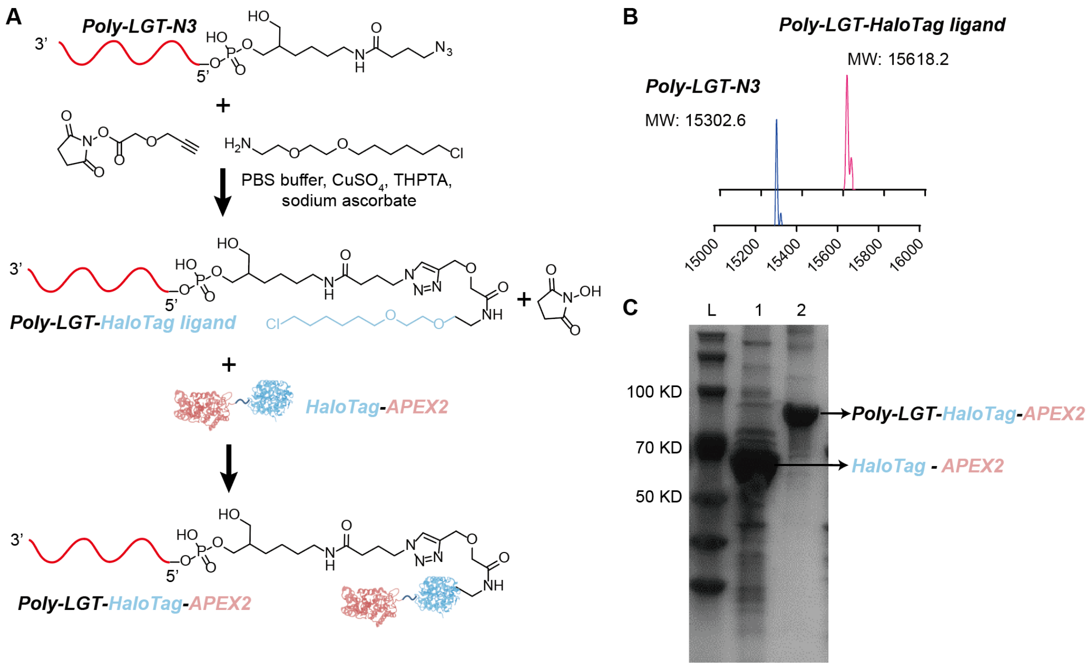


**Figure S3. Characterization of origami-APEX2 system. A)** Schematic diagram of ssDNA conjugated with HaloTag-APEX2 in origami-APEX2 system. **B)** Mass spectrum of ssDNA-$N_3$ and ssDNA-HaloTag ligand. **C)** SDS-PAGE characterization of conjugation of ssDNA-HaloTag ligand with HaloTag-APEX2.

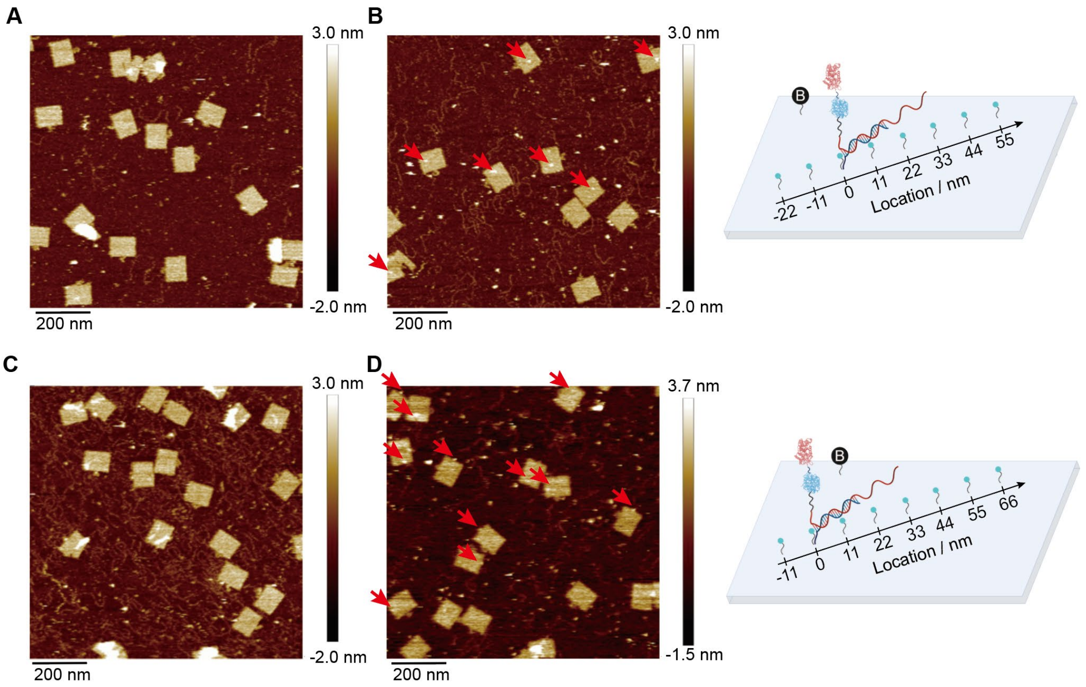


**Figure S4. AFM image of origami - APEX2 conjugations. A)** origami (135-AC). **B)** origami (135-AC) + APEX2. **C)** origami (161-AC). **D)** origami (161-AC) + APEX2.

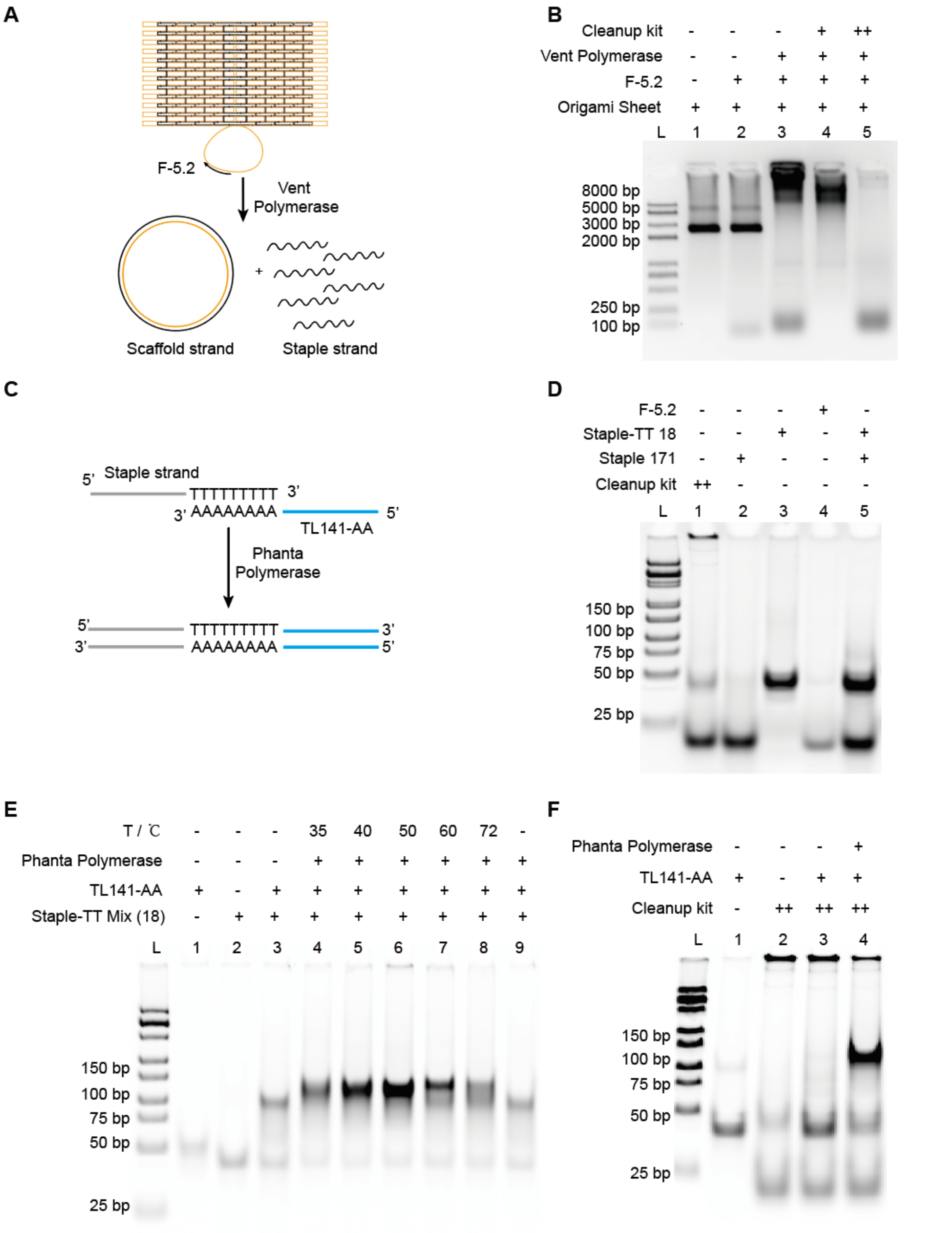


**Figure S5. Validation of workflow of APEX2-DNA origami system. A)** Schematic diagram illustrating the separation of staple strand from origami by Vent polymerase. **B)** Agarose gel analysis for characterizing the separation of staple strand from origami. "Cleanup kit +" and "Cleanup kit ++" are samples recovered from the first and second columns during the purification process, respectively. **C)** Schematic diagram illustrating the extension of staple strand with primer TL141-AA. **D)** 10% native PAGE characterization of the isolated staple strand. **E)** Optimization of temperature in staple stand extension. **F)** Polymerize single-stranded DNA into double-stranded by Phanta polymerase.

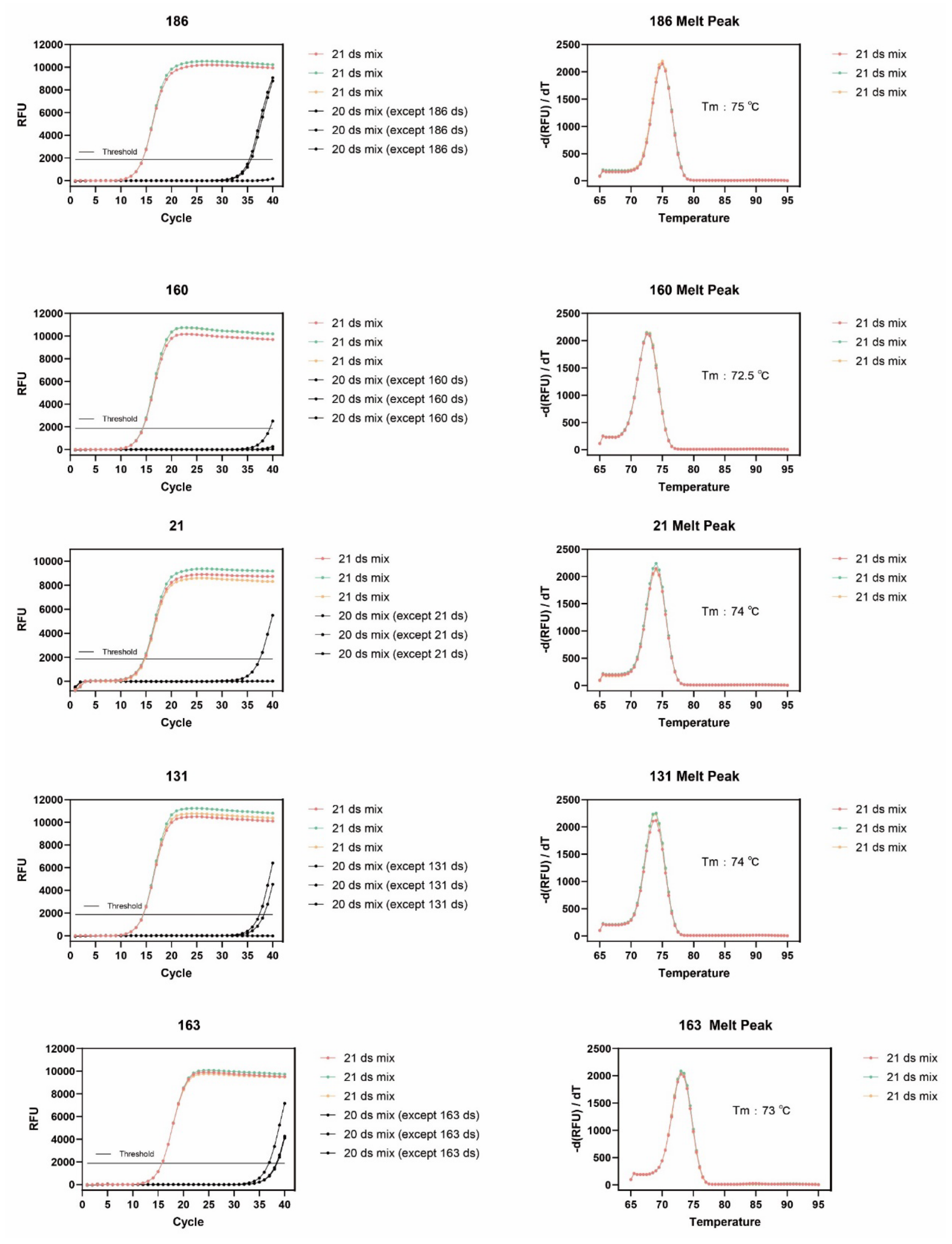


**Figure S6.** Verification of primer specificity.

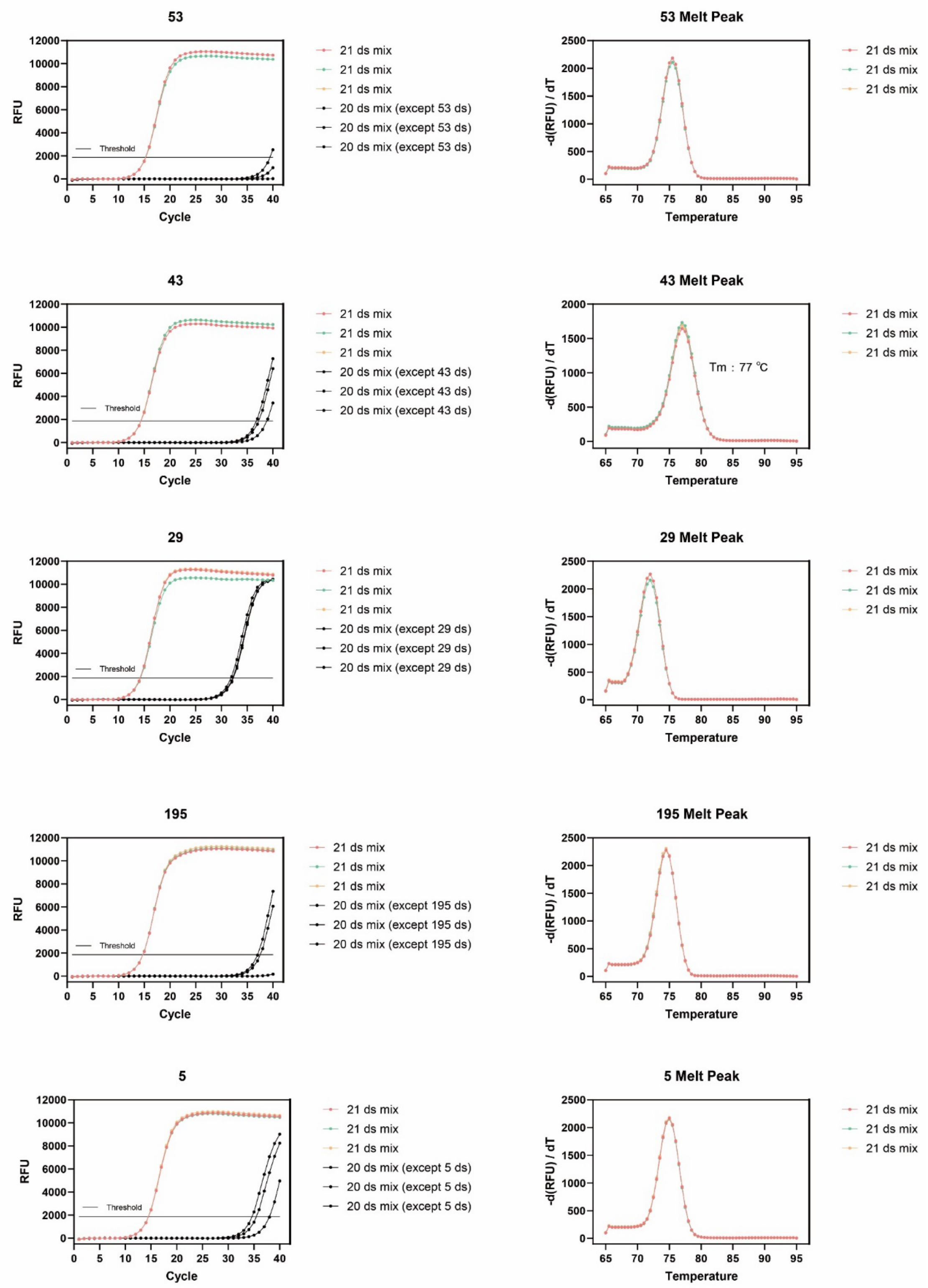


**Figure S6.** Verification of primer specificity.

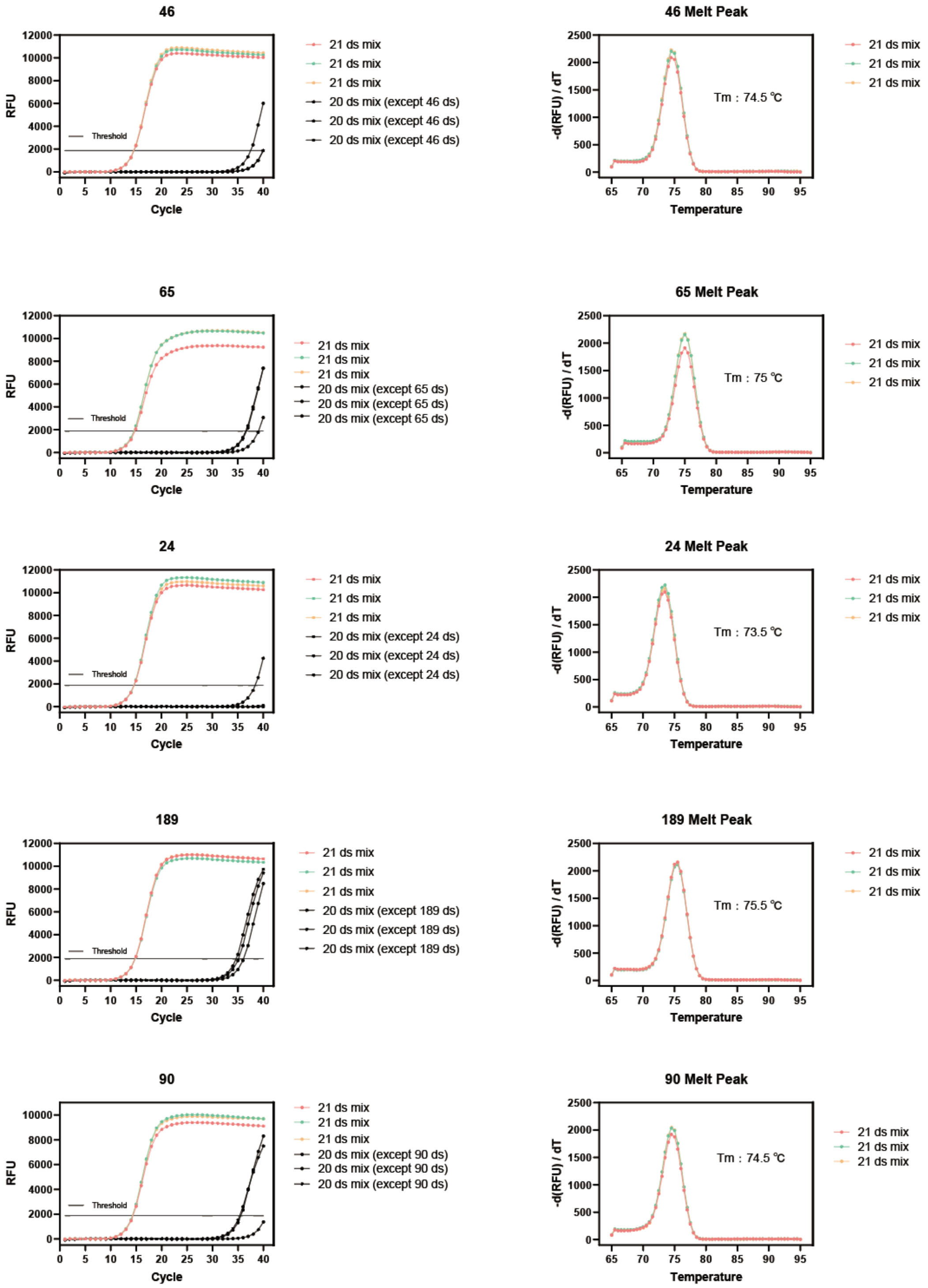


**Figure S6.** Verification of primer specificity.

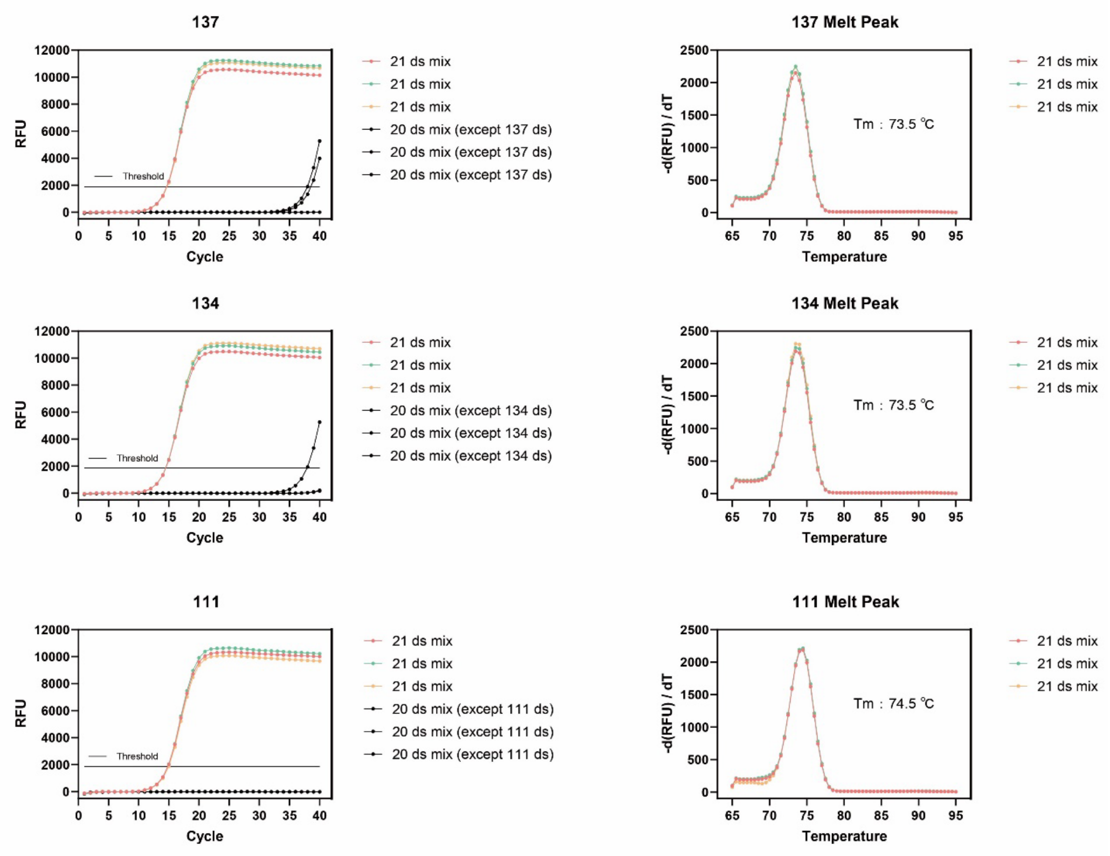


**Figure S6.** Verification of primer specificity.

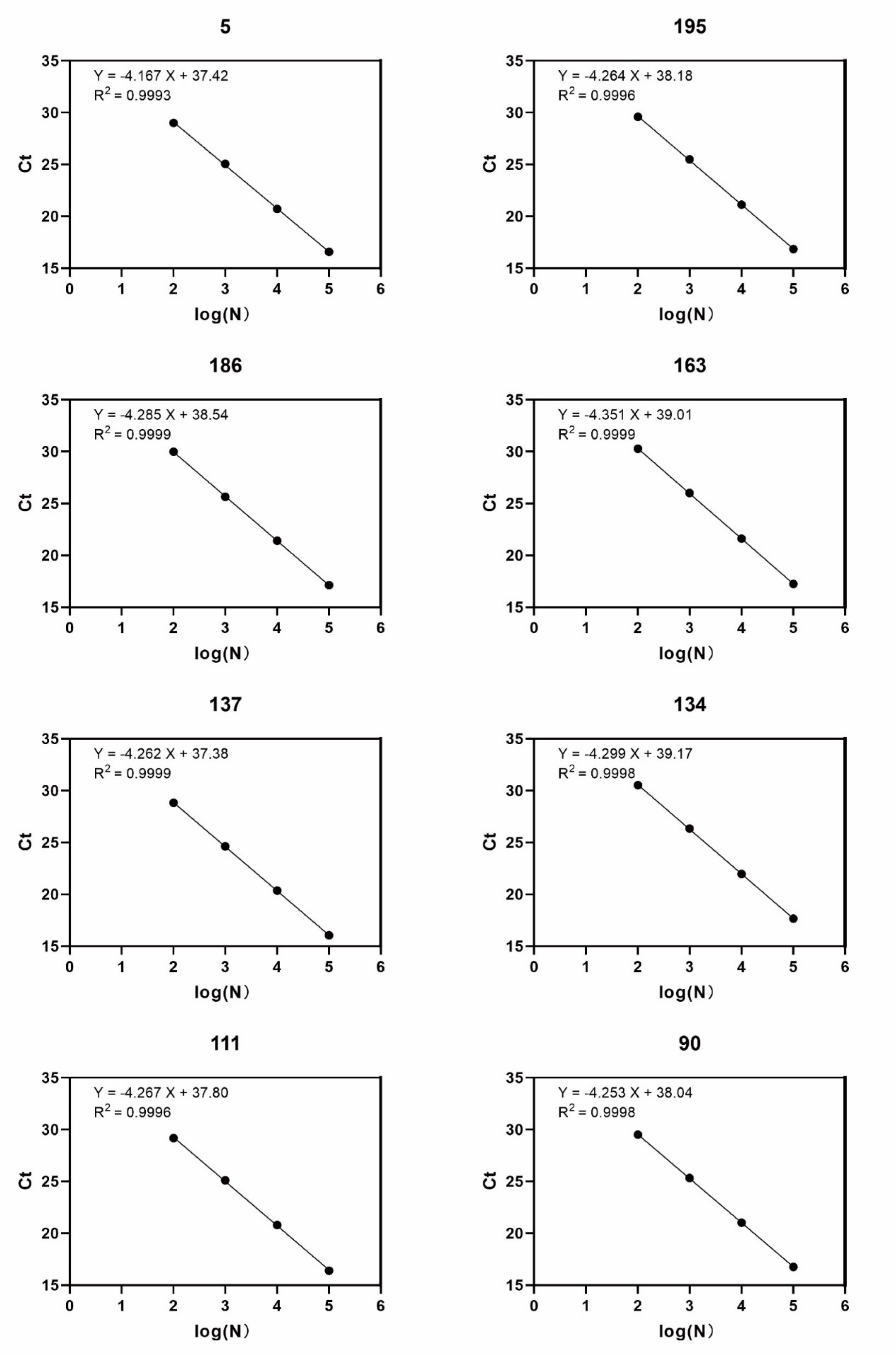


**Figure S7.** A simple linear regression curve of the Ct value and the logarithm of the dsDNA number was used to further obtain the amplification efficiency.

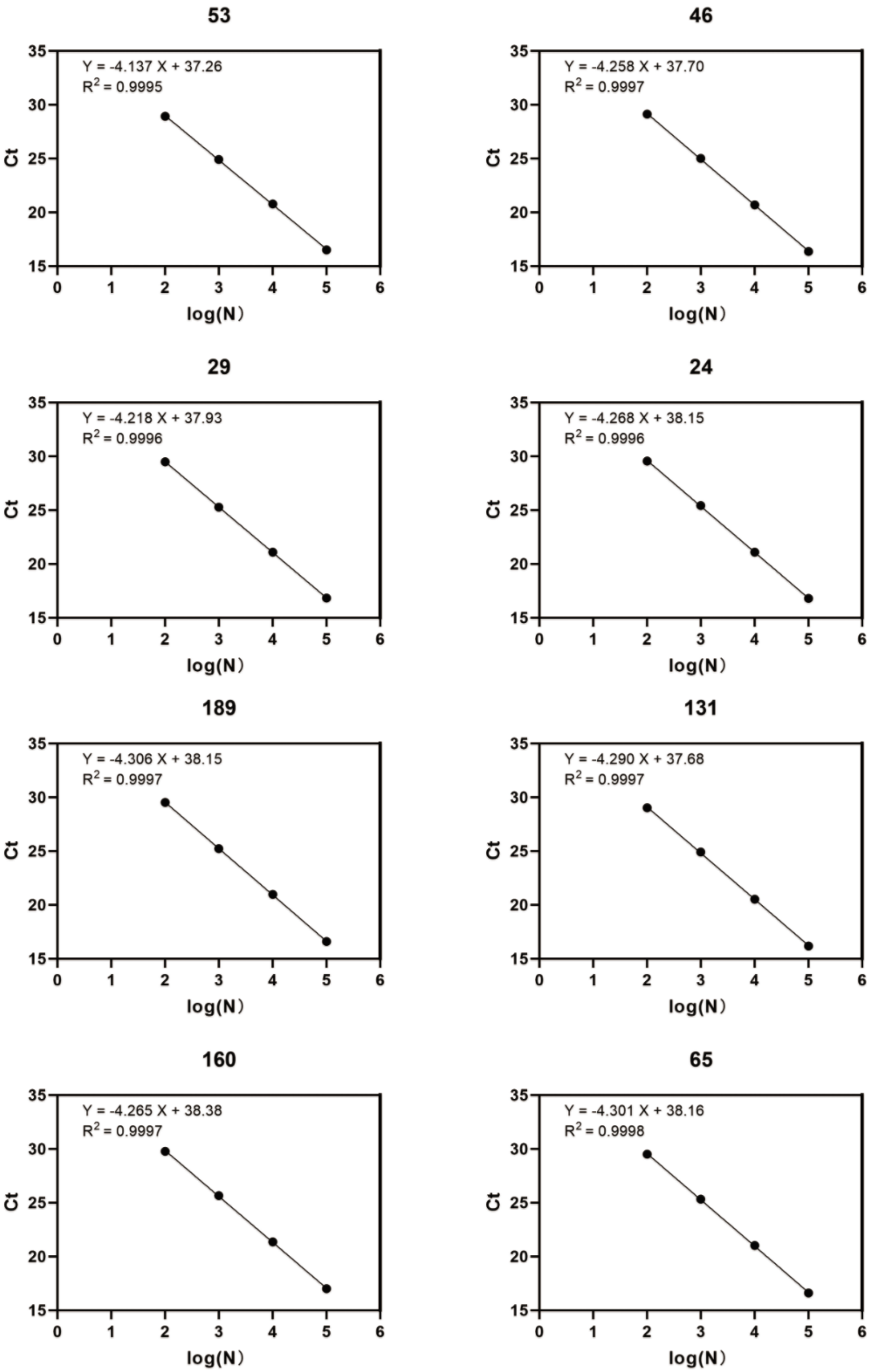


**Figure S7.** A simple linear regression curve of the Ct value and the logarithm of the dsDNA number was used to further obtain the amplification efficiency.

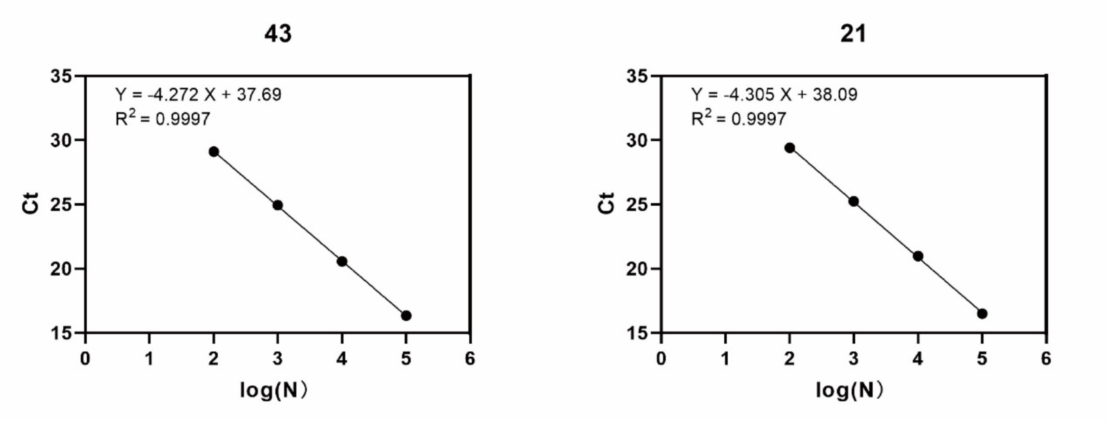


**Figure S7**. A simple linear regression curve of the Ct value and the logarithm of the dsDNA number was used to further obtain the amplification efficiency.

**Table S1.** Calculation of amplification efficiency.

| Staple | Slope (k) | E (%) |
|---|---|---|
| 90 | -4.253 | 71.84 |
| 65 | -4.301 | 70.81 |
| 46 | -4.258 | 71.73 |
| 24 | -4.268 | 71.51 |
| 43 | -4.272 | 71.43 |
| 21 | -4.305 | 70.72 |
| 131 | -4.29 | 71.04 |
| 111 | -4.267 | 71.54 |
| 134 | -4.299 | 70.85 |
| 137 | -4.262 | 71.65 |
| 160 | -4.265 | 71.58 |
| 163 | -4.351 | 69.76 |
| 186 | -4.285 | 71.15 |
| 189 | -4.306 | 70.7 |
| 195 | -4.264 | 71.6 |
| 53 | -4.137 | 74.47 |
| 29 | -4.218 | 72.62 |
| 5 | -4.167 | 73.77 |

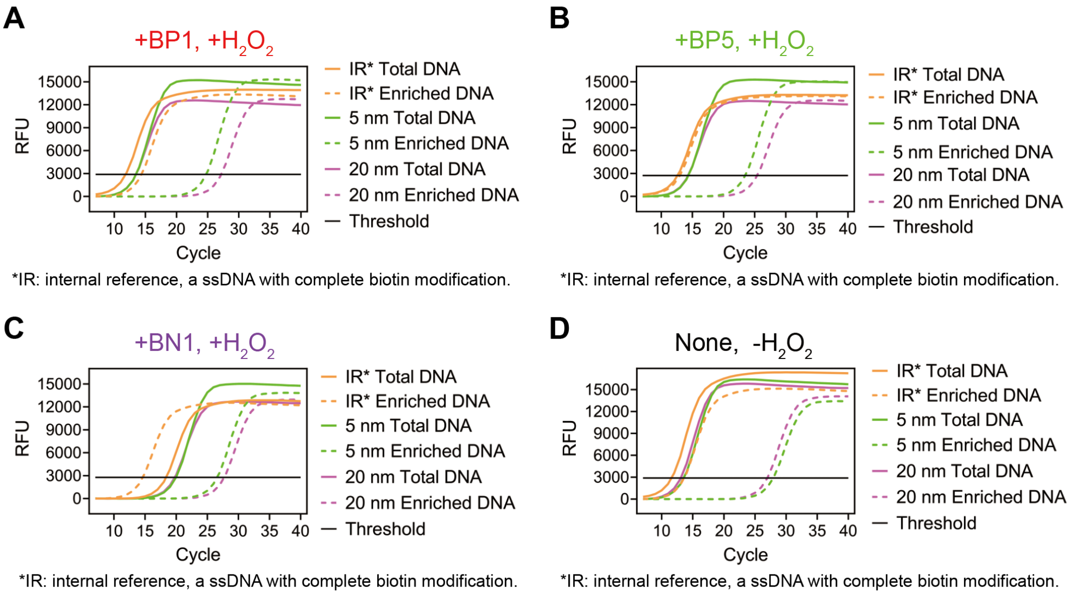


**Figure S8.** Representative qPCR curves for quantifying labeling efficiencies in Figure 2F. **A)** + BP1, + $H_2O_2$. **B)** + BP5, + $H_2O_2$. **C)** + BN1, + $H_2O_2$. **D)** + None, **-** $H_2O_2$.

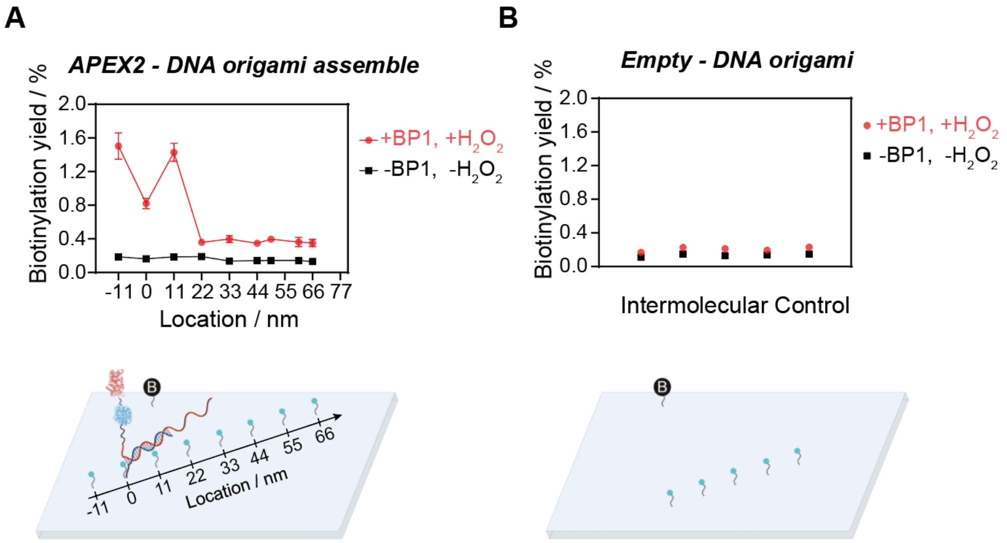


**Figure S9.** Plot of biotinylation yield against phenol location in APEX2-DNA origami assemble **(A)** and empty- DNA origami **(B)**.

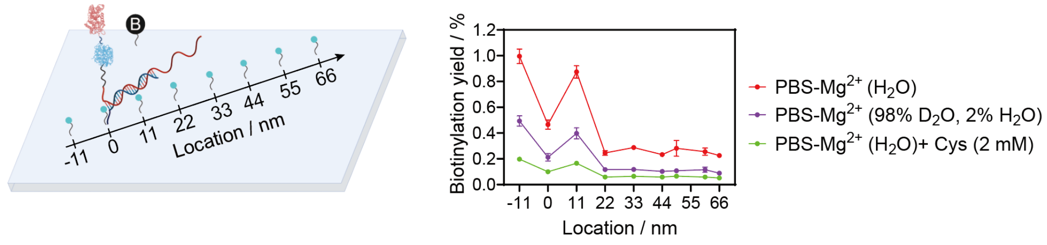


**Figure S10.** Effect of Cys and $D_2O$ on labeling efficiencies of APEX2.

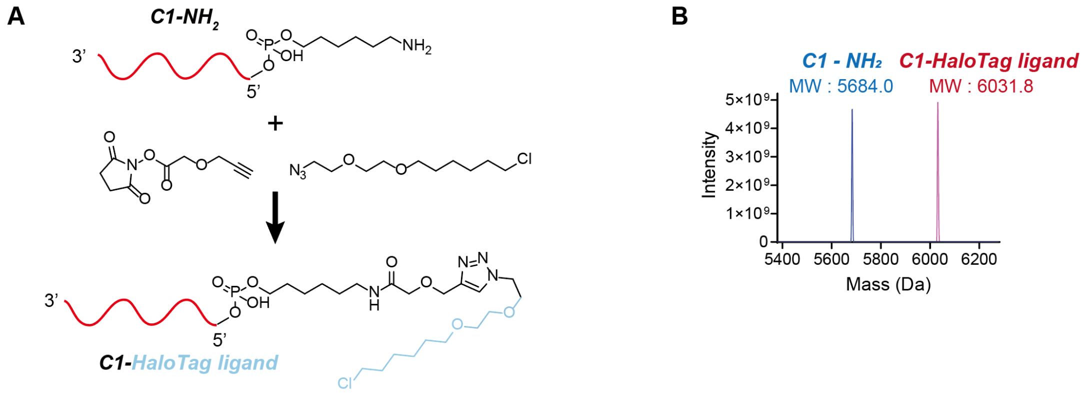


**Figure S11. Characterization of TurboID (APEX2) - dsDNA system. A)** Schematic diagram of ssDNA conjugated with HaloTag ligand. **B)** Mass spectrum of ssDNA-$NH_2$ and ssDNA-HaloTag ligand.

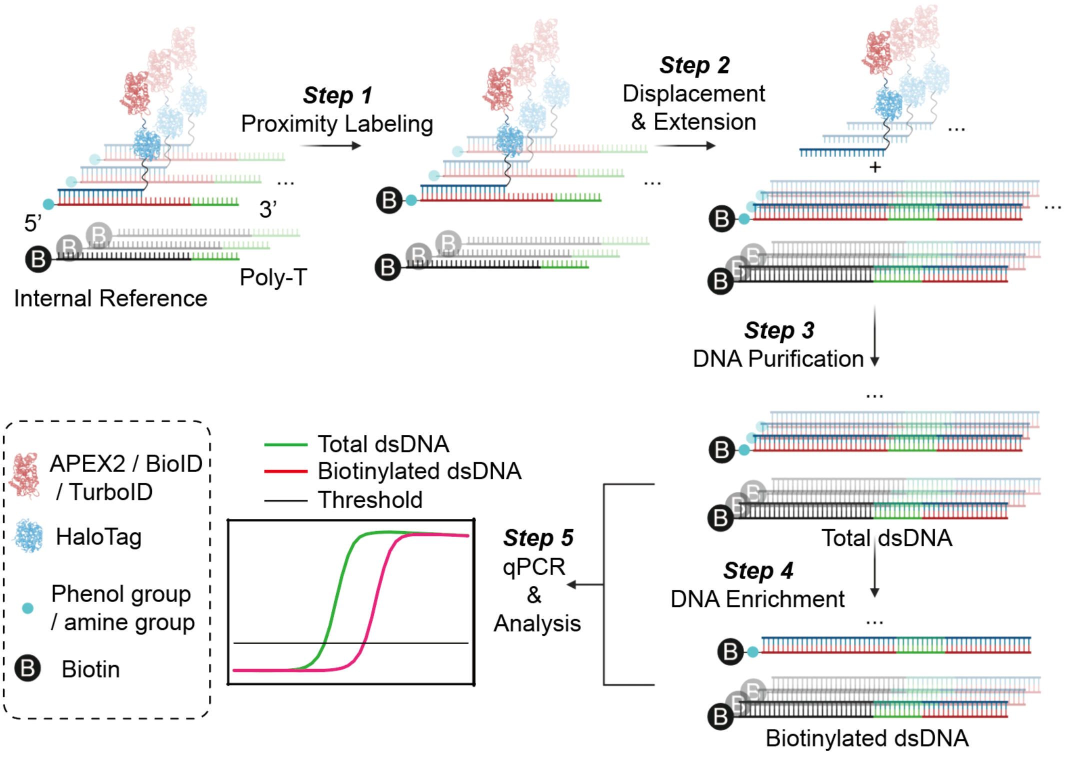


**Figure S12.** Workflow of the dsDNA-labeling-enriching-qPCR system.

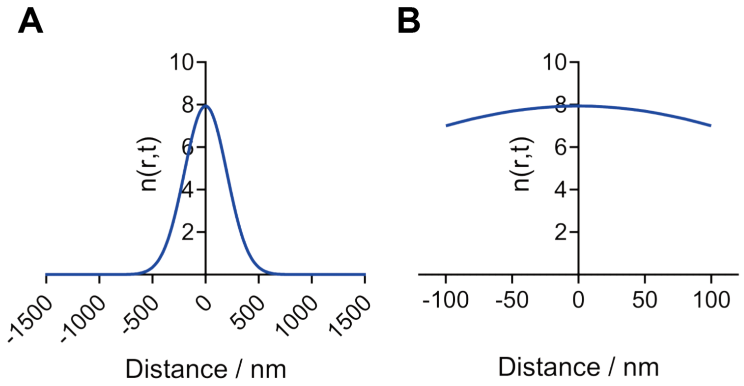


**Figure S13. The simulation of APEX2 functioning through diffusive labeling mechanism. A)** The simulated concentration gradient of phenoxyl radicals as a function of distance, D = 200 $\mu m^2/s$, t= 0.1 ms. **B)** The enlarged distance-dependent concentration gradient of phenoxyl radicals within 100 nm.

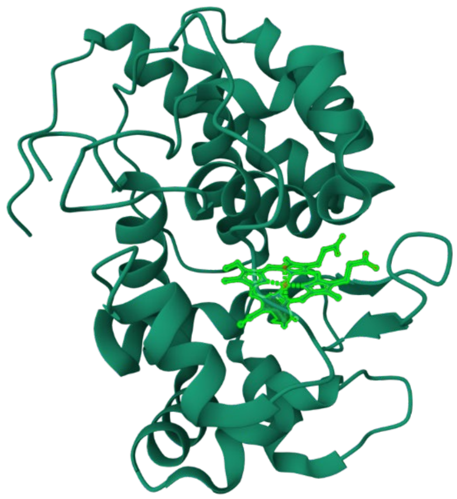

**Figure S14. The Crystal Structures of Ascorbate Peroxidase (APX, PDB ID: 1OAF).** APEX2 has a similar structure, with the active center Heme located in a solvent-exposed position.

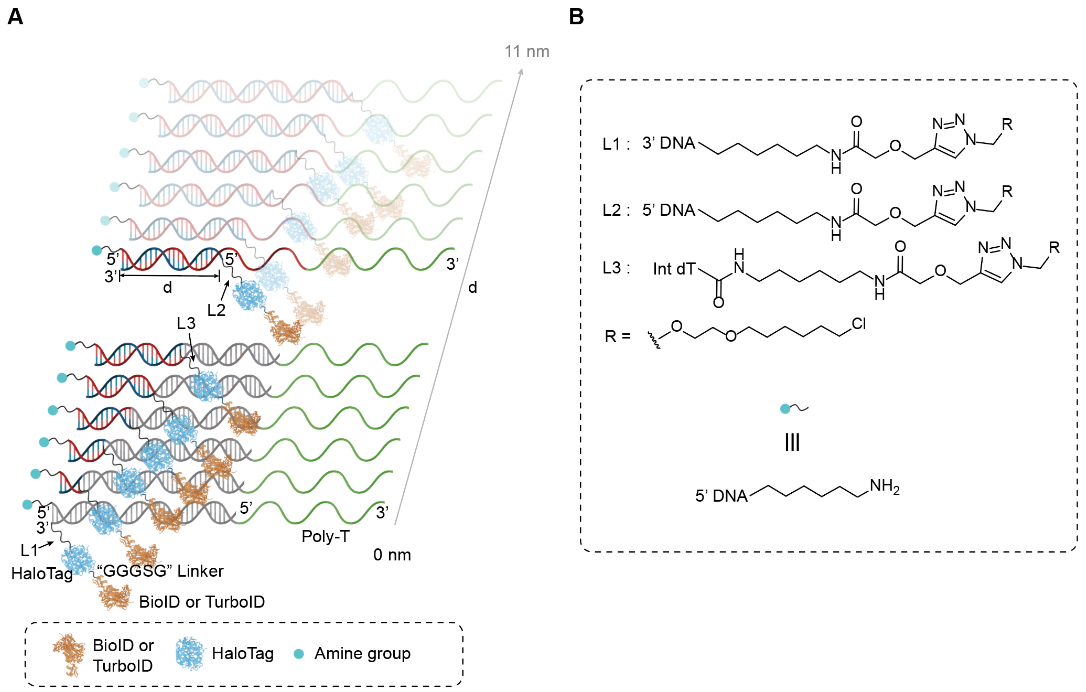


**Figure S15. A)** Schematic diagram of different distance of TurboID with amine group. **B)** Chemical structure of linkers and molecules in **A**.

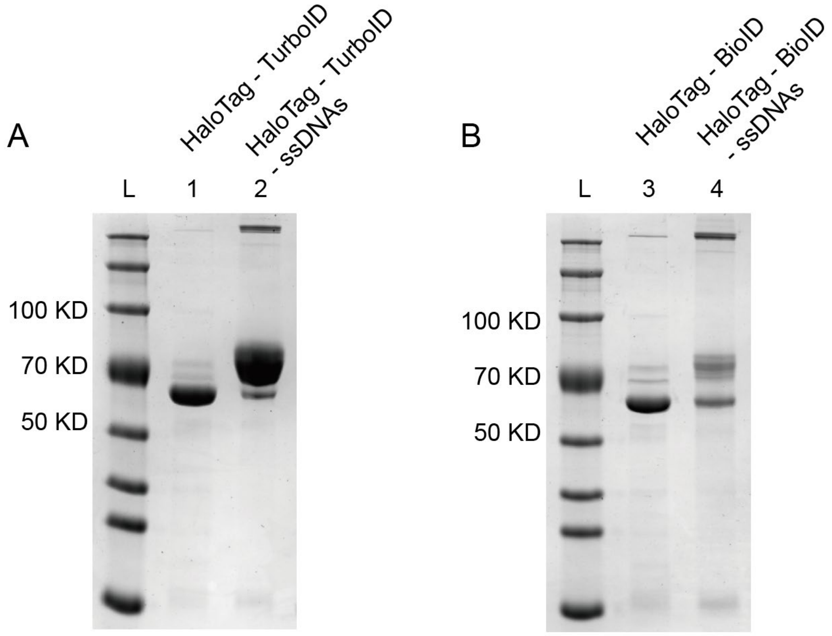


**Figure S16. A)** SDS-PAGE characterization of conjugation of ssDNAs-HaloTag ligand with HaloTag-TurboID. **B)** SDS-PAGE characterization of conjugation of ssDNAs-HaloTag ligand with HaloTag -BioID.

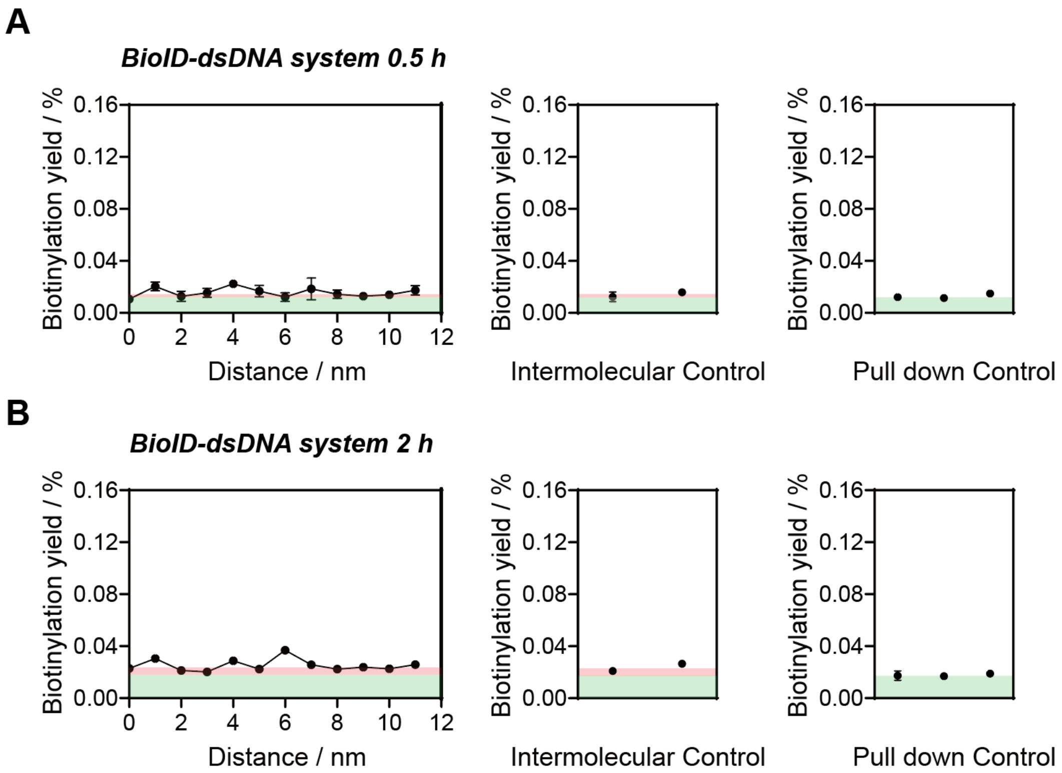


**Figure S17. A)** Labeling efficiencies of BioID to amines with different distances spaced by dsDNA in 0.5 h. **B)** Labeling efficiencies of BioID to amines with different distances spaced by dsDNA in 2 h. Pink areas represent intermolecular labeling background and green areas represent pull down background.

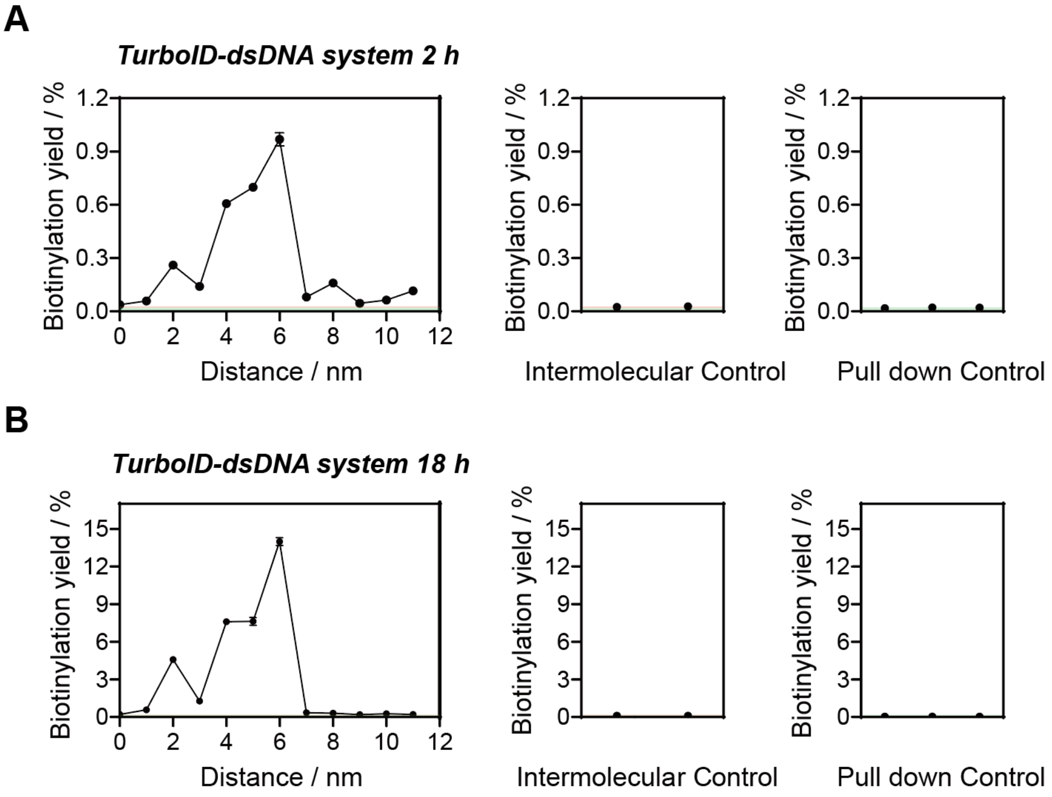


**Figure S18. A)** Labeling efficiencies of TurboID to amines with different distances spaced by dsDNA in 2 h. **B)** Labeling efficiencies of TurboID to amines with different distances spaced by dsDNA in 18 h. Pink areas represent intermolecular labeling background and green areas represent pull down background.

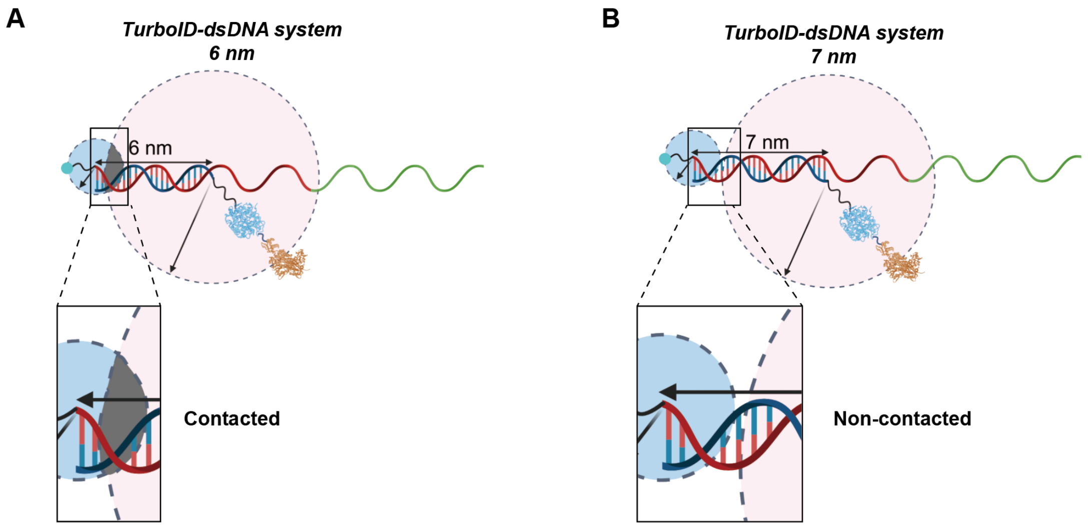


**Figure S19. A)** Schematic diagram of the contact-dependent labeling model of TurboID. **B)** Schematic diagram of the Non-contact-dependent labeling model of TurboID.

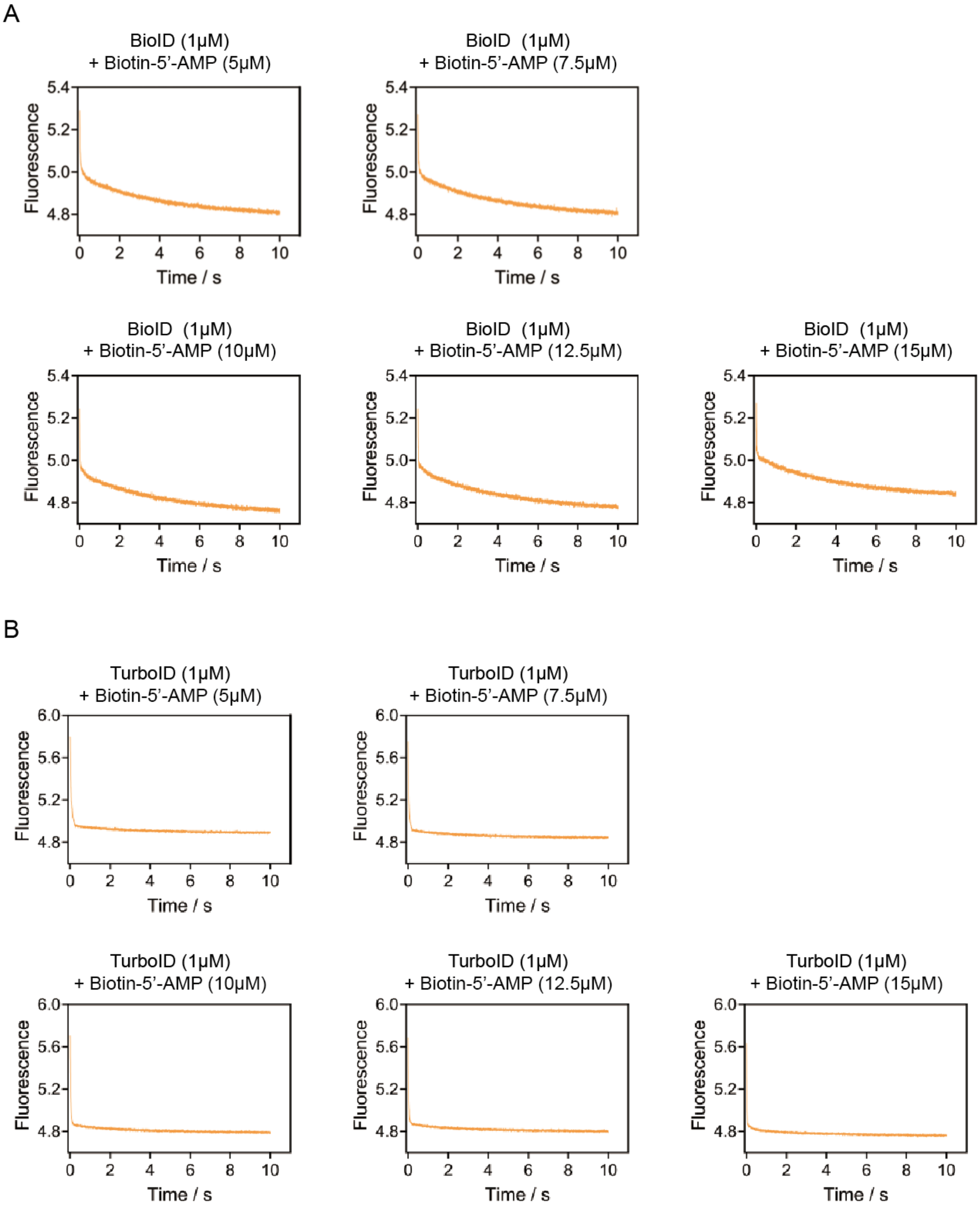


**Figure S20.** Bimolecular association of BioID (A) and TurboID (B) with biotin-5’-AMP.

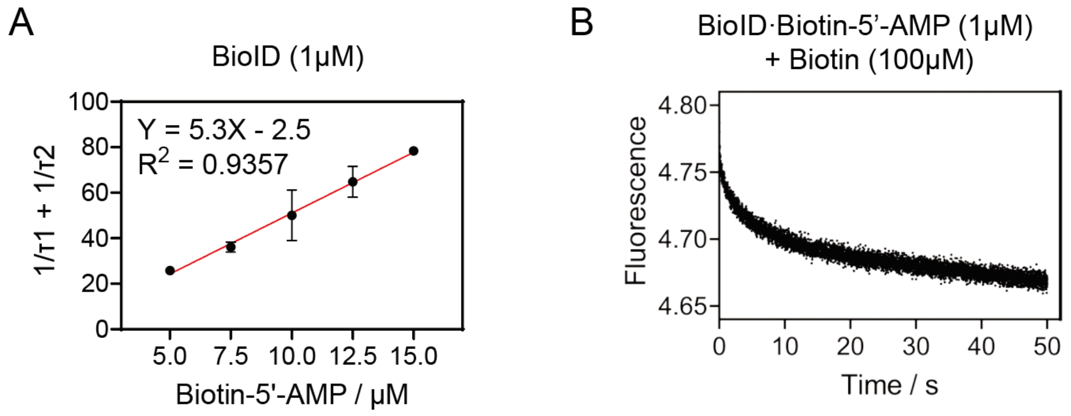


**Figure S21. A)** Dependence of the sum of the apparent rates (1/τ1 and 1/τ2) of the two kinetic phases on ligand concentration for BioID. **B)** Dissociation of the complexes of BioID bound to biotin-5'-AMP measured by stopped-flow fluorescence.

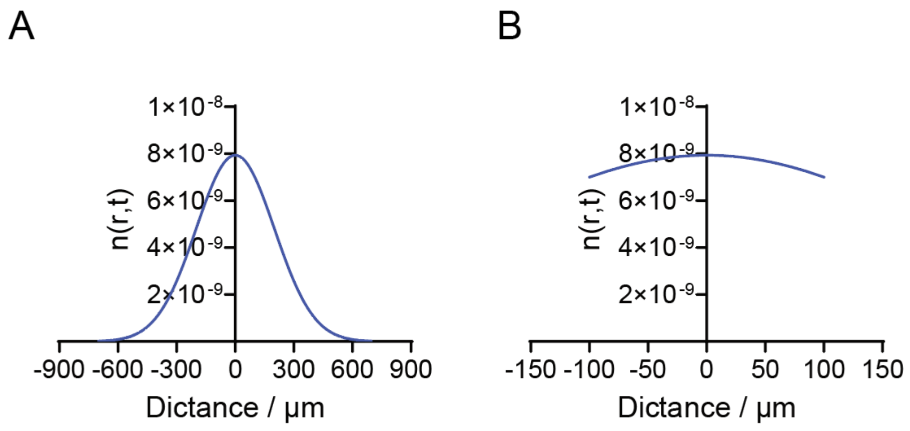


**Figure S22. A)** Simulated biotin-5'-AMP concentration gradient as a function of distance, D = 200 $\mu m^2/s$, t= 100 s. **B)** The enlarged distance-dependent biotin-5'-AMP concentration gradient within 100 µm.

**Table S2.** Sequence of specific staple strands used in Figure 2E.

| Location | Name | Sequence 5'-3' |
|---|---|---|
| 55 nm | 90 (5'-$NH_2$) | $NH_2$-TGCTCAGTCAGTCTCTGAATTTACCAGGAGGT TTTTTTTTTTTTTTTTTTTTTTTTT |
| 49.5 nm | 65 (5'-$NH_2$) | $NH_2$-TAAGCGTCGAAGGATTAGGATTAGTACCGCCA TTTTTTTTTTTTTTTTTTTTTTTTT |
| 33 nm | 46 (5'-$NH_2$) | $NH_2$- GACTTGAGAGACAAAAGGGCGACAAGTTACCA TTTTTTTTTTTTTTTTTTTTTTTTT |
| 22 nm | 24 (5'-$NH_2$) | $NH_2$-AAAAGTAATATCTTACCGAAGCCCAACACTAT TTTTTTTTTTTTTTTTTTTTTTTTT |
| 16.5 nm | 131 (5'-$NH_2$) | $NH_2$-CATAACCCGAGGCATAGTAAGAGCTTTTTAAG TTTTTTTTTTTTTTTTTTTTTTTTT |
| 11 nm | 111 (5'-$NH_2$) | $NH_2$- AAGAGGAACGAGCTTCAAAGCGAAGATACATT TTTTTTTTTTTTTTTTTTTTTTTTT |
| 5.5 nm | 134 (5'-$NH_2$) | $NH_2$- GAAGCAAAAAAGCGGATTGCATCAGATAAAAA TTTTTTTTTTTTTTTTTTTTTTTTT |
| 0 nm | 137 (5'-$NH_2$) | $NH_2$-CGAGTAGAACTAATAGTAGTAGCAAACCCTCA TTTTTTTTTTTTTTTTTTTTTTTTT |
| -5.5 nm | 160 (5'-$NH_2$) | $NH_2$-CAATAAATACAGTTGATTCCCAATTTAGAGAG TTTTTTTTTTTTTTTTTTTTTTTTT |
| -11 nm | 163 (5'-$NH_2$) | $NH_2$-CAACGCAATTTTTGAGAGATCTACTGATAATC TTTTTTTTTTTTTTTTTTTTTTTTT |
| -16.5 nm | 186 (5'-$NH_2$) | $NH_2$- TCAGGTCACTTTTGCGGGAGAAGCAGAATTAG TTTTTTTTTTTTTTTTTTTTTTTTT |
| -22 nm | 189 (5'-$NH_2$) | $NH_2$-CATGTCAAGATTCTCCGTGGGAACCGTTGGTG TTTTTTTTTTTTTTTTTTTTTTTTT |
| Internal Reference | 195 (5'-Biotin) | Biotin- TGGTTTTTAACGTCAAAGGGCGAAGAACCATC TTTTTTTTTTTTTTTTTTTTTTTTT |
| APEX2 anchoring strand (handle strand) | 135-AC | TCAGAAGCCTCCAACAGGTCAGGATCTGCGAA tttACACACACACACACACAC |

**Table S3.** Sequence of specific staple strands used in Figure S8.

| Location | Name | Sequence 5'-3' |
|---|---|---|
| 66 nm | 90 (5'-$NH_2$) | $NH_2$-TGCTCAGTCAGTCTCTGAATTTACCAGGAGGT TTTTTTTTTTTTTTTTTTTTTTTTT |
| 60.5 nm | 65 (5'-$NH_2$) | $NH_2$-TAAGCGTCGAAGGATTAGGATTAGTACCGCCA TTTTTTTTTTTTTTTTTTTTTTTTT |
| 49.5 nm | 43 (5'-$NH_2$) | $NH_2$-GTTTGCCACCTCAGAGCCGCCACCGATACAGG TTTTTTTTTTTTTTTTTTTTTTTTT |
| 44 nm | 46 (5'-$NH_2$) | $NH_2$-GACTTGAGAGACAAAAGGGCGACAAGTTACCA TTTTTTTTTTTTTTTTTTTTTTTTT |
| 33 nm | 24 (5'-$NH_2$) | $NH_2$-AAAAGTAATATCTTACCGAAGCCCAACACTAT TTTTTTTTTTTTTTTTTTTTTTTTT |
| 22 nm | 111 (5'-$NH_2$) | $NH_2$-AAGAGGAACGAGCTTCAAAGCGAAGATACATT TTTTTTTTTTTTTTTTTTTTTTTTT |
| 11 nm | 137 (5'-$NH_2$) | $NH_2$-CGAGTAGAACTAATAGTAGTAGCAAACCCTCA TTTTTTTTTTTTTTTTTTTTTTTTT |
| 0 nm | 163 (5'-$NH_2$) | $NH_2$-CAACGCAATTTTTGAGAGATCTACTGATAATC TTTTTTTTTTTTTTTTTTTTTTTTT |
| -11 nm | 189 (5'-$NH_2$) | $NH_2$-CATGTCAAGATTCTCCGTGGGAACCGTTGGTG TTTTTTTTTTTTTTTTTTTTTTTTT |
| Internal Reference | 195 (5'-Biotin) | $NH_2$-TGGTTTTTAACGTCAAAGGGCGAAGAACCATC TTTTTTTTTTTTTTTTTTTTTTTTT |
| APEX2 anchoring strand (handle strand) | 161-AC | TCCATATACATACAGGCAAGGCAACTTTATTT tttACACACACACACACACACAC |
| Control | 21 (5'-$NH_2$) | $NH_2$-ATTGAGGGTAAAGGTGAATTATCAATCACCGG TTTTTTTTTTTTTTTTTTTTTTTTT |
| Control | 131 (5'-$NH_2$) | $NH_2$-CATAACCCGAGGCATAGTAAGAGCTTTTTAAG TTTTTTTTTTTTTTTTTTTTTTTTT |
| Control | 134 (5'-$NH_2$) | $NH_2$-GAAGCAAAAAAGCGGATTGCATCAGATAAAAA TTTTTTTTTTTTTTTTTTTTTTTTT |
| Control | 160 (5'-$NH_2$) | $NH_2$-CAATAAATACAGTTGATTCCCAATTTAGAGAG TTTTTTTTTTTTTTTTTTTTTTTTT |
| Control | 186 (5'-$NH_2$) | $NH_2$-TCAGGTCACTTTTGCGGGAGAAGCAGAATTAG TTTTTTTTTTTTTTTTTTTTTTTTT |

**Table S4.** Specific primers for qPCR.

| Name | Sequence 5’-3’ |
|---|---|
| P-46 | GACTTGAGAGACAAAAGGGCGACAAG |
| P-24 | AAAAGTAATATCTTACCGAAGCCCAACAC |
| P-111 | AAGAGGAACGAGCTTCAAAGCG |
| P-163 | CAACGCAATTTTTGAGAGATCTACTGATAATC |
| P-189 | CATGTCAAGATTCTCCGTGGGAACC |
| P-195 | TGGTTTTTAACGTCAAAGGGCGAAG |
| P-137 | CGAGTAGAACTAATAGTAGTAGCAAACCC |
| P-90 | TGCTCAGTCAGTCTCTGAATTTACCAGG |
| P-65 | TAAGCGTCGAAGGATTAGGATTAGTACCG |
| P-43 | GTTTGCCACCTCAGAGCCG |
| P-21 | ATTGAGGGTAAAGGTGAATTATCAATCACCG |
| P-131 | CATAACCCGAGGCATAGTAAGAGC |
| P-134 | GAAGCAAAAAAGCGGATTGCATCAG |
| P-160 | CAATAAATACAGTTGATTCCCAATTTAGAGAG |
| P-186 | TCAGGTCACTTTTGCGGGAGAAG |
| P-53 | CAAGCAAGACGCGCCTG |
| P-29 | GGTATTAAGAACAAGAAAAATAATTAAAGCCA |
| P-5 | ATCGGCTGCGAGCATGTAG |
| P141 | GTCTACTATATGAGTCTACTAGCCTTATGATACTCTG |

**Table S5.** Primers for assembly and workflow.

| Name | Sequence 5’-3’ |
|---|---|
| Detecting strand-Cy5 | CACTCCTAGTATCACAACGAAGC-Cy5 |
| Poly-LGT-$N_3$ (anti-handle strand) | GCTTCGTTGTGATACTAGGAGTGTTGTGTGTGTGTGTGTGTG TGTTTT -$N_3$ |
| Poly-LGT-$NH_2$ (anti-handle strand) | GCTTCGTTGTGATACTAGGAGTGTTGTGTGTGTGTGTGTGTG TGTTTT -$NH_2$ |
| F-5.2 | AGTGAGGCCACCGAGTAAAAGAGTCTGTCCATCACGC |
| TL 141-AA (Adapter strand) | GTCTACTATATGAGTCTACTAGCCTTATGATACTCTGATaaaaaaa aaaaaaaaaaaaa |

**Table S6.** Sequence of ssDNA used in dsDNA scaffold system (Figure 3D).

| Location | Name | Sequence 5'-3' |
|---|---|---|
| 0 nm | 163-C (0$NH_2$) | GATTATCAGTAGATCTCTCAAAAATTGCGTTG-$NH_2$ |
| 2 nm | 160-C (7$NH_2$) | CTCTCTAAATTGGGAATCAACTGTA/i$NH_2$C6dT/TTATTG |
| 4 nm | 137-C (13$NH_2$) | TGAGGGTTTGCTACTACTA/i$NH_2$C6dT/TAGTTCTACTCG |
| 6 nm | C1 (5-$NH_2$) | $NH_2$-ACTATGCCTCGGGTTATG |
| 7 nm | C2 (5-$NH_2$) | $NH_2$-CTTCGGTAAGATATTACTTTT |
| 8 nm | C3 (5-$NH_2$) | $NH_2$-TGATAATTCACCTTTACCCTCAAT |
| 9 nm | C4 (5-$NH_2$) | $NH_2$-ACTTGTCGCCCTTTGTCTCTCAAGTC |
| 10 nm | C5 (5-$NH_2$) | $NH_2$-TGTATCGGTGGCGGCTCTGAGGTGGCAAAC |
| 11 nm | C6 (5-$NH_2$) | $NH_2$-ATGGCGGTACTAATCCTAATCCTTCGACGCTTA |
| 0 nm | 163 (5'-$NH_2$) | $NH_2$-CAACGCAATTTTTGAGAGATCTACTGATAATC TTTTTTTTTTTTTTTTTTTTTTTTT |
| 2 nm | 160 (5'-$NH_2$) | $NH_2$-CAATAAATACAGTTGATTCCCAATTTAGAGAG TTTTTTTTTTTTTTTTTTTTTTTTT |
| 4 nm | 137 (5'-$NH_2$) | $NH_2$-CGAGTAGAACTAATAGTAGTAGCAAACCCTCA TTTTTTTTTTTTTTTTTTTTTTTTT |
| 6 nm | 131 (5'-$NH_2$) | $NH_2$-CATAACCCGAGGCATAGTAAGAGCTTTTTAAG TTTTTTTTTTTTTTTTTTTTTTTTT |
| 7 nm | 24 (5'-$NH_2$) | $NH_2$-AAAAGTAATATCTTACCGAAGCCCAACACTAT TTTTTTTTTTTTTTTTTTTTTTTTT |
| 8 nm | 21 (5'-$NH_2$) | $NH_2$-ATTGAGGGTAAAGGTGAATTATCAATCACCGG TTTTTTTTTTTTTTTTTTTTTTTTT |
| 9 nm | 46 (5'-$NH_2$) | $NH_2$-GACTTGAGAGACAAAAGGGCGACAAGTTACCA TTTTTTTTTTTTTTTTTTTTTTTTT |
| 10 nm | 43 (5'-$NH_2$) | $NH_2$-GTTTGCCACCTCAGAGCCGCCACCGATACAGG TTTTTTTTTTTTTTTTTTTTTTTTT |
| 11 nm | 65 (5'-$NH_2$) | $NH_2$-TAAGCGTCGAAGGATTAGGATTAGTACCGCCA TTTTTTTTTTTTTTTTTTTTTTTTT |
| Control | 111 (5'-$NH_2$) | $NH_2$-AAGAGGAACGAGCTTCAAAGCGAAGATACATT TTTTTTTTTTTTTTTTTTTTTTTTT |
| Control | 134 (5'-$NH_2$) | $NH_2$-GAAGCAAAAAAGCGGATTGCATCAGATAAAAA TTTTTTTTTTTTTTTTTTTTTTTTT |
| Internal Reference | 195 (5'-Biotin) | $NH_2$-TGGTTTTTAACGTCAAAGGGCGAAGAACCATC TTTTTTTTTTTTTTTTTTTTTTTTT |

Scaffold strands: 163-C (0$NH_2$), 160-C (7$NH_2$), 137-C (13$NH_2$), C1 (5-$NH_2$), C2 (5-$NH_2$), C3 (5-$NH_2$), C4 (5-$NH_2$), C5 (5-$NH_2$), C6 (5-$NH_2$).

**Table S7.** Sequence of specific staple strands used in dsDNA scaffold system (Figure 4B, C, Figure S15, S16).

| Location | Name | Sequence 5'-3' |
|---|---|---|
| 0 nm | 163-C (0$NH_2$) | GATTATCAGTAGATCTCTCAAAAATTGCGTTG-$NH_2$ |
| 1 nm | 111-C (4-NH2) | AATGTATCTTCGCTTTGAAGCTCGTTCC/iNH2C6dT/CTT |
| 2 nm | 160-C (7$NH_2$) | CTCTCTAAATTGGGAATCAACTGTA/i$NH_2$C6dT/TTATTG |
| 3 nm | 189-C (10NH2) | CACCAACGGTTCCCACGGAGAA/iNH2C6dT/CTTGACATG |
| 4 nm | 137-C (13$NH_2$) | TGAGGGTTTGCTACTACTA/i$NH_2$C6dT/TAGTTCTACTCG |
| 5 nm | 134-C (16-NH2) | TTTTTATCTGATGCAA/iNH2C6dT/CCGCTTTTTTGCTTC |
| 6 nm | C1 (5-$NH_2$) | $NH_2$-ACTATGCCTCGGGTTATG |
| 7 nm | C2 (5-$NH_2$) | $NH_2$-CTTCGGTAAGATATTACTTTT |
| 8 nm | C3 (5-$NH_2$) | $NH_2$-TGATAATTCACCTTTACCCTCAAT |
| 9 nm | C4 (5-$NH_2$) | $NH_2$-ACTTGTCGCCCTTTTGTCTCTCAAGTC |
| 10 nm | C5 (5-$NH_2$) | $NH_2$-TGTATCGGTGGCGGCTCTGAGGTGGCAAAC |
| 11 nm | C6 (5-$NH_2$) | $NH_2$-ATGGCGGTACTAATCCTAATCCTTCGACGCTTA |
| 0 nm | 163 (5'-$NH_2$) | $NH_2$-CAACGCAATTTTTGAGAGATCTACTGATAATC<br>TTTTTTTTTTTTTTTTTTTTTTTTT |
| 1 nm | 111 (5'-NH2) | $NH_2$-AAGAGGAACGAGCTTCAAAGCGAAGATACATT<br>TTTTTTTTTTTTTTTTTTTTTTTTT |
| 2 nm | 160 (5'-$NH_2$) | $NH_2$-CAATAAATACAGTTGATTCCCAATTTAGAGAG<br>TTTTTTTTTTTTTTTTTTTTTTTTT |
| 3 nm | 189 (5'-NH2) | $NH_2$-CATGTCAAGATTCTCCGTGGGAACCGTTGGTG<br>TTTTTTTTTTTTTTTTTTTTTTTTT |
| 4 nm | 137 (5'-$NH_2$) | $NH_2$-CGAGTAGAACTAATAGTAGTAGCAAACCCTCA<br>TTTTTTTTTTTTTTTTTTTTTTTTT |
| 5 nm | 134 (5'-NH2) | $NH_2$-GAAGCAAAAAAGCGGATTGCATCAGATAAAAA<br>TTTTTTTTTTTTTTTTTTTTTTTTT |
| 6 nm | 131 (5'-$NH_2$) | $NH_2$-CATAACCCGAGGCATAGTAAGAGCTTTTTAAG<br>TTTTTTTTTTTTTTTTTTTTTTTTT |
| 7 nm | 24 (5'-$NH_2$) | $NH_2$-AAAAGTAATATCTTACCGAAGCCCAACACTAT<br>TTTTTTTTTTTTTTTTTTTTTTTTT |
| 8 nm | 21 (5'-$NH_2$) | $NH_2$-ATTGAGGGTAAAGGTGAATTATCAATCACCGG<br>TTTTTTTTTTTTTTTTTTTTTTTTT |
| 9 nm | 46 (5'-$NH_2$) | $NH_2$-GACTTGAGAGACAAAAGGGCGACAAGTTACCA<br>TTTTTTTTTTTTTTTTTTTTTTTTT |
| 10 nm | 43 (5'-$NH_2$) | $NH_2$-GTTTGCCACCTCAGAGCCGCCACCGATACAGG<br>TTTTTTTTTTTTTTTTTTTTTTTTT |
| 11 nm | 65 (5'-$NH_2$) | $NH_2$-TAAGCGTCGAAGGATTAGGATTAGTACCGCCA<br>TTTTTTTTTTTTTTTTTTTTTTTTT |
| Control | 90 (5'-$NH_2$) | $NH_2$-TGCTCAGTCAGTCTCTGAATTTACCAGGAGGT<br>TTTTTTTTTTTTTTTTTTTTTTTTT |
| Control | 186 (5'-$NH_2$) | $NH_2$-TCAGGTCACTTTTGCGGGAGAAGCAGAATTAG<br>TTTTTTTTTTTTTTTTTTTTTTTTT |

| | | |
|---|---|---|
| Control | 53 - TT | $NH_2$-CAAGCAAGACGCGCCTGTTTATCAAGAATCGC TTTTTTTTTTTTTTTTTTTTTTTTTT |
| Control | 29 - TT | $NH_2$-GGTATTAAGAACAAGAAAAATAATTAAAGCCA TTTTTTTTTTTTTTTTTTTTTTTTTT |
| Control | 5 - TT | $NH_2$-ATCGGCTGCGAGCATGTAGAAACCAGCTATAT TTTTTTTTTTTTTTTTTTTTTTTTTT |
| Internal Reference | 195 (5'-Biotin) | Biotin-TGGTTTTTAACGTCAAAGGGCGAAGAACCATC TTTTTTTTTTTTTTTTTTTTTTTTTT |

Scaffold strands: 163-C (0$NH_2$), 111-C (4-$NH_2$), 160-C (7$NH_2$), 189-C (10$NH_2$), 137-C (13$NH_2$), 134-C (16-$NH_2$), C1 (5-$NH_2$), C2 (5-$NH_2$), C3 (5-$NH_2$), C4 (5-$NH_2$), C5 (5-$NH_2$), C6 (5-$NH_2$).

**Table S8.** Sequence of staple strands used in Rectangular DNA origami.

| Name | Sequence 5’-3’ |
|---|---|
| 1 | CAAGCCCAATAGGAACCCATGTACCGTAACAC |
| 2 | TCTTACCAGCCAGTTACAAAATAAATGAAATA |
| 3 | CCTAATTTACGCTAACGAGCGTCTATATCGCG |
| 4 | CTAATTTATCTTTCCTTATCATTCATCCTGAA |
| 5 | ATCGGCTGCGAGCATGTAGAAACCAGCTATAT |
| 6 | AATTACTACAAATTCTTACCAGTAATCCCATC |
| 7 | GCGTTATAGAAAAAGCCTGTTTAGAAGGCCGG |
| 8 | TAGAATCCCTGAGAAGAGTCAATAGGAATCAT |
| 9 | TTAAGACGTTGAAAACATAGCGATTTAAATCA |
| 10 | TTTAACGTTCGGGAGAAACAATAATTTTCCCT |
| 11 | CTTTTACACAGATGAATATACAGTAAGCGCCA |
| 12 | GGATTTAGCGTATTAAATCCTTTGTTTTCAGG |
| 13 | CGACAACTAAGTATTAGACTTTACAGCCGGAA |
| 14 | TAGCCCTACCAGCAGAAGATAAAAACATTTGA |
| 15 | ACGAACCAAAACATCGCCATTAAATGGTGGTT |
| 16 | CGGCCTTGCTGGTAATATCCAGAACGAACTGA |
| 17 | TGCCTTGACTGCCTATTTCGGAACAGGGATAG |
| 18 | AATGCCCCGTAACAGTGCCCGTATGTGAATTT |
| 19 | AACCAGAGACCCTCAGAACCGCCAGGGGTCAG |
| 20 | GAGCCGCCCCACCACCGGAACCGCCTAAAACA |
| 21 | ATTGAGGGTAAAGGTGAATTATCAATCACCGG |
| 22 | TTATTCATAGGGAAGGTAAATATTCATTCAGT |
| 23 | GCAATAGCGCAGATAGCCGAACAATTCAACCG |
| 24 | AAAAGTAATATCTTACCGAAGCCCAACACTAT |
| 25 | CTCAGAGCCACCACCCTCATTTTCCTATTATT |
| 26 | TATTTTGCTCCCAATCCAAATAAGTGAGTTAA |
| 27 | ATTATTTAACCCAGCTACAATTTTCAAGAACG |
| 28 | TAAGTCCTACCAAGTACCGCACTCTTAGTTGC |
| 29 | GGTATTAAGAACAAGAAAAATAATTAAAGCCA |
| 30 | AGGCGTTACAGTAGGGCTTAATTGACAATAGA |
| 31 | ACGCTCAAAATAAGAATAAACACCGTGAATTT |
| 32 | CTGTAAATCATAGGTCTGAGAGACGATAAATA |
| 33 | ATCAAAATCGTCGCTATTAATTAACGGATTCG |
| 34 | ACAGAAATCTTTGAATACCAAGTTCCTTGCTT |
| 35 | CCTGATTGAAAGAAATTGCGTAGACCCGAACG |
| 36 | AGATTAGATTTAAAAGTTTGAGTACACGTAAA |
| 37 | TTATTAATGCCGTCAATAGATAATCAGAGGTG |
| 38 | GAATGGCTAGTATTAACACCGCCTCAACTAAT |
| 39 | AGGCGGTCATTAGTCTTTAATGCGCAATATTA |
| 40 | CCGCCAGCCATTGCAACAGGAAAAATATTTTT |

| | |
|---|---|
| 41 | AGTGTACTTGAAAGTATTAAGAGGCCGCCACC |
| 42 | CTGAAACAGGTAATAAGTTTTAACCCCTCAGA |
| 43 | GTTTGCCACCTCAGAGCCGCCACCGATACAGG |
| 44 | GCCACCACTCTTTTCATAATCAAACCGTCACC |
| 45 | AGCGCCAACCATTTGGGAATTAGATTATTAGC |
| 46 | GACTTGAGAGACAAAAGGGCGACAAGTTACCA |
| 47 | GCCCAATACCGAGGAAACGCAATAGGTTTACC |
| 48 | GAAGGAAAATAAGAGCAAGAAACAACAGCCAT |
| 49 | CCCTCAGAACCGCCACCCTCAGAACTGAGACT |
| 50 | AGGTTTTGAACGTCAAAAATGAAAGCGCTAAT |
| 51 | TTTTGTTTAAGCCTTAAATCAAGAATCGAGAA |
| 52 | AATGCAGACCGTTTTTATTTTCATCTTGCGGG |
| 53 | CAAGCAAGACGCGCCTGTTTATCAAGAATCGC |
| 54 | AATGGTTTACAACGCCAACATGTAGTTCAGCT |
| 55 | CATATTTAGAAATACCGACCGTGTTACCTTTT |
| 56 | AAATCAATGGCTTAGGTTGGGTTACTAAATTT |
| 57 | TAACCTCCATATGTGAGTGAATAAACAAAATC |
| 58 | AACCTACCGCGAATTATTCATTTCCAGTACAT |
| 59 | GCGCAGAGATATCAAAATTATTTGACATTATC |
| 60 | CTAAAATAGAACAAAGAAACCACCAGGGTTAG |
| 61 | ATTTTGCGTCTTTAGGAGCACTAAGCAACAGT |
| 62 | GCGTAAGAGAGAGCCAGCAGCAAAAAGGTTAT |
| 63 | GCCACGCTATACGTGGCACAGACAACGCTCAT |
| 64 | GGAAATACCTACATTTTGACGCTCACCTGAAA |
| 65 | TAAGCGTCGAAGGATTAGGATTAGTACCGCCA |
| 66 | CCTCAAGAATACATGGCTTTTGATAGAACCAC |
| 67 | TCGGCATTCCGCCGCCAGCATTGACGTTCCAG |
| 68 | CACCAGAGTTCGGTCATAGCCCCCGCCAGCAA |
| 69 | TCACAATCGTAGCACCATTACCATCGTTTTCA |
| 70 | AATCACCAAATAGAAAATTCATATATAACGGA |
| 71 | ATCAGAGAAAGAACTGGCATGATTTTATTTTG |
| 72 | ATACCCAAGATAACCCACAAGAATAAACGATT |
| 73 | TATCACCGTACTCAGGAGGTTTAGCGGGGTTT |
| 74 | GAGGCGTTAGAGAATAACATAAAAGAACACCC |
| 75 | CTTTACAGTTAGCGAACCTCCCGACGTAGGAA |
| 76 | CCAGACGAGCGCCCAATAGCAAGCAAGAACGC |
| 77 | TCATTACCCGACAATAAACAACATATTTAGGC |
| 78 | TTTTAGTTTTTCGAGCCAGTAATAAATTCTGT |
| 79 | AGAGGCATAATTTCATCTTCTGACTATAACTA |
| 80 | TTGAATTATGCTGATGCAAATCCACAAATATA |
| 81 | TATGTAAACCTTTTTTAATGGAAAAATTACCT |
| 82 | TGGATTATGAAGATGATGAAACAAAATTTCAT |

| | |
|---|---|
| 83 | GAGCAAAAACTTCTGAATAATGGAAGAAGGAG |
| 84 | ATCAACAGTCATCATATTCCTGATTGATTGTT |
| 85 | CGGAATTATTGAAAGGAATTGAGGTGAAAAAT |
| 86 | GCCAACAGTCACCTTGCTGAACCTGTTGGCAA |
| 87 | CTAAAGCAAGATAGAACCCTTCTGAATCGTCT |
| 88 | GAAATGGATTATTTACATTGGCAGACATTCTG |
| 89 | GGAAAGCGACCAGGCGGATAAGTGAATAGGTG |
| 90 | TGCTCAGTCAGTCTCTGAATTTACCAGGAGGT |
| 91 | TGCCTTTAGTCAGACGATTGGCCTGCCAGAAT |
| 92 | TGAGGCAGGCGTCAGACTGTAGCGTAGCAAGG |
| 93 | ACGCAAAGGTCACCAATGAAACCAATCAAGTT |
| 94 | CCGGAAACACACCACGGAATAAGTAAGACTCC |
| 95 | TGAACAAACAGTATGTTAGCAAACTAAAAGAA |
| 96 | TTATTACGGTCAGAGGGTAATTGAATAGCAGC |
| 109 | TGAGTTTCGTCACCAGTACAAACTTAATTGTA |
| 110 | TTTTAATTGCCCGAAAGACTTCAATTCCAGAG |
| 111 | AAGAGGAACGAGCTTCAAAGCGAAGATACATT |
| 112 | TTTCATTTGGTCAATAACCTGTTTAATCAATA |
| 113 | TCGCAAATGGGGCGCGAGCTGAAATAATGTGT |
| 114 | AGACAGTCATTCAAAAGGGTGAGATATCATAT |
| 115 | AGGTAAAGAAATCACCATCAATATAATATTTT |
| 116 | GCTCATTTTCGCATTAAATTTTTGAGCTTAGA |
| 117 | GTTAAAATTTTAACCAATAGGAACCCGGCACC |
| 118 | TTCGCCATTGCCGGAAACCAGGCAAACAGTAC |
| 119 | GCTTCTGGTCAGGCTGCGCAACTGTGTTATCC |
| 120 | GCATAAAGTTCCACACAACATACGAAACAATT |
| 121 | GCTCACAATGTAAAGCCTGGGGTGGGTTTGCC |
| 122 | CCGAAATCCGAAAATCCTGTTTGAAATACCGA |
| 123 | CCAGCAGGGGCAAAATCCCTTATAAAGCCGGC |
| 124 | GAACGTGGCGAGAAAGGAAGGGAACAAACTAT |
| 125 | CTTAAACATCAGCTTGCTTTCGAGAAACAGTT |
| 126 | TCGGTTTAGCTTGATACCGATAGTCCAACCTA |
| 127 | CTCATCTTGAGGCAAAAGAATACACTCCCTCA |
| 128 | AAACGAAATGACCCCCAGCGATTATTCATTAC |
| 129 | GAATAAGGACGTAACAAAGCTGCTGACGGAAA |
| 130 | CCAAATCACTTGCCCTGACGAGAACGCCAAAA |
| 131 | CATAACCCGAGGCATAGTAAGAGCTTTTTAAG |
| 132 | GGAATTACTCGTTTACCAGACGACAAAAGATT |
| 133 | TGTAGCATTCCACAGACAGCCCTCATCTCCAA |
| 134 | GAAGCAAAAAAGCGGATTGCATCAGATAAAAA |
| 135 | TCAGAAGCCTCCAACAGGTCAGGATCTGCGAA |
| 136 | TCAATTCTTTTAGTTTGACCATTACCAGACCG |

| | |
|---|---|
| 137 | CGAGTAGAACTAATAGTAGTAGCAAACCCTCA |
| 138 | ACCGTTCTAAATGCAATGCCTGAGAGGTGGCA |
| 139 | TATATTTTAGCTGATAAATTAATGTTGTATAA |
| 140 | AAATAATTTTAAATTGTAAACGTTGATATTCA |
| 141 | GCAAATATCGCGTCTGGCCTTCCTGGCCTCAG |
| 142 | GGCGATCGCACTCCAGCCAGCTTTGCCATCAA |
| 143 | GAAGATCGGTGCGGGCCTCTTCGCAATCATGG |
| 144 | GTGAGCTAGTTTCCTGTGTGAAATTTGGGAAG |
| 145 | TCATAGCTACTCACATTAATTGCGCCCTGAGA |
| 146 | GAATAGCCGCAAGCGGTCCACGCTCCTAATGA |
| 147 | GAGTTGCACGAGATAGGGTTGAGTAAGGGAGC |
| 148 | CCCCGATTTAGAGCTTGACGGGGAAATCAAAA |
| 149 | CAATGACACTCCAAAAGGAGCCTTACAACGCC |
| 150 | AAAAAAGGACAACCATCGCCCACGCGGGTAAA |
| 151 | GCGAAACATGCCACTACGAAGGCATGCGCCGA |
| 152 | ATACGTAAAAGTACAACGGAGATTTCATCAAG |
| 153 | ACGAGTAGTGACAAGAACCGGATATACCAAGC |
| 154 | AGTAATCTTAAATTGGGCTTGAGAGAATACCA |
| 155 | CCAAAATATAATGCAGATACATAAACACCAGA |
| 156 | CATTCAACGCGAGAGGCTTTTGCATATTATAG |
| 157 | CGTAACGATCTAAAGTTTTGTCGTGAATTGCG |
| 158 | TACCTTTAAGGTCTTTACCCTGACAAAGAAGT |
| 159 | CAAAAATCATTGCTCCTTTTGATAAGTTTCAT |
| 160 | CAATAAATACAGTTGATTCCCAATTTAGAGAG |
| 161 | TCCATATACATACAGGCAAGGCAACTTTATTT |
| 162 | GGTAGCTAGGATAAAAATTTTTAGTTAACATC |
| 163 | CAACGCAATTTTTGAGAGATCTACTGATAATC |
| 164 | CTTTCATCCCCAAAAACAGGAAGACCGGAGAG |
| 165 | AGAAAAGCAACATTAAATGTGAGCATCTGCCA |
| 166 | CAGCTGGCGGACGACGACAGTATCGTAGCCAG |
| 167 | GTTTGAGGGAAAGGGGGATGTGCTAGAGGATC |
| 168 | ACTGCCCGCCGAGCTCGAATTCGTTATTACGC |
| 169 | CCCGGGTACTTTCCAGTCGGGAAACGGGCAAC |
| 170 | AGTTTGGAGCCCTTCACCGCCTGGTTGCGCTC |
| 171 | AGCTGATTACAAGAGTCCACTATTGAGGTGCC |
| 172 | GTAAAGCACTAAATCGGAACCCTAGTTGTTCC |
| 173 | ATATATTCTTTTTTCACGTTGAAAATAGTTAG |
| 174 | AATAATAAGGTCGCTGAGGCTTGCAAAGACTT |
| 175 | CGCCTGATGGAAGTTTCCATTAAACATAACCG |
| 176 | TTTCATGAAAATTGTGTCGAAATCTGTACAGA |
| 177 | TTTCAACTATAGGCTGGCTGACCTTGTATCAT |
| 178 | CCAGGCGCTTAATCATTGTGAATTACAGGTAG |

| | |
|---|---|
| 179 | TTTGCCAGATCAGTTGAGATTTAGTGGTTTAA |
| 180 | AAAGATTCAGGGGGTAATAGTAAACCATAAAT |
| 181 | ACGTTAGTAAATGAATTTTCTGTAAGCGGAGT |
| 182 | TTTTTGCGCAGAAAACGAGAATGAATGTTTAG |
| 183 | AAACAGTTGATGGCTTAGAGCTTATTTAAATA |
| 184 | CAAAATTAAAGTACGGTGTCTGGAAGAGGTCA |
| 185 | TGCAACTAAGCAATAAAGCCTCAGTTATGACC |
| 186 | TCAGGTCACTTTTGCGGGAGAAGCAGAATTAG |
| 187 | CTGTAATATTGCCTGAGAGTCTGGAAAACTAG |
| 188 | ACCCGTCGTCATATGTACCCCGGTAAAGGCTA |
| 189 | CATGTCAAGATTCTCCGTGGGAACCGTTGGTG |
| 190 | ATTAAGTTCGCATCGTAACCGTGCGAGTAACA |
| 191 | TAGATGGGGGGTAACGCCAGGGTTGTGCCAAG |
| 192 | GCCAGCTGCCTGCAGGTCGACTCTGCAAGGCG |
| 193 | CTTGCATGCATTAATGAATCGGCCCGCCAGGG |
| 194 | TGGACTCCCTTTTCACCAGTGAGACCTGTCGT |
| 195 | TGGTTTTTAACGTCAAAGGGCGAAGAACCATC |
| 196 | ACCCAAATCAAGTTTTTTGGGGTCAAAGAACG |
| 197 | AAAGGCCGAAAGGAACAACTAAAGCTTTCCAG |
| 198 | GAGAATAGCTTTTGCGGGATCGTCGGGTAGCA |
| 199 | GCTCCATGAGAGGCTTTGAGGACTAGGGAGTT |
| 200 | ACGGCTACTTACTTAGCCGGAACGCTGACCAA |
| 201 | CGATTTTAGAGGACAGATGAACGGCGCGACCT |
| 202 | CTTTGAAAAGAACTGGCTCATTATTTAATAAA |
| 203 | ACTGGATAACGGAACAACATTATTACCTTATG |
| 204 | ACGAACTAGCGTCCAATACTGCGGAATGCTTT |

**Protein sequences**

HaloTag-APEX2:
MGSEIGTGFPFDPHYVEVLGERMHYVDVGPRDGTPVLFLHGNPTSSYVWRNIIPHVA
PTHRCIAPDLIGMGKSDKPDLGYFFDDHVRFMDAFIEALGLEEVVLVIHDWGSALGF
HWAKRNPERVKGIAFMEFIRPIPTWDEWPEFARETFQAFRTTDVGRKLIIDQNVFIEGT
LPMGVVRPLTEVEMDHYREPFLNPVDREPLWRFPNELPIAGEPANIVALVEEYMDWL
HQSPVPKLLFWGTPGVLIPPAEAARLAKSLPNCKAVDIGPGLNLLQEDNPDLIGSEIAR
WLSTLEISGGGGSGASGKSYPTVSADYQDAVEKAKKKLRGFIAEKRCAPLMLRLAFH
SAGTFDKGTKTGGPFGTIKHPAELAHSANNGLDIAVRLLEPLKAEFPILSYADFYQLAG
VVAVEVTGGPKVPFHPGREDKPEPPPEGRLPDPTKGSDHLRDVFGKAMGLTDQDIVA
LSGGHTIGAAHKERSGFEGPWTSNPLIFDNSYFTELLSGEKEGLLQLPSDKALLSDPVF
RPLVDKYAADEDAFFADYAEAHQKLSELGFADAGSDYKDDDDKGLNDIFEAQKIEW
HEHHHHHH
His-tag, Flag-tag, AviTag, nhe1 cleavage site, HaloTag, APEX2, GS linker

HaloTag-TurboID:
MGSEIGTGFPFDPHYVEVLGERMHYVDVGPRDGTPVLFLHGNPTSSYVWRNIIPHVA
PTHRCIAPDLIGMGKSDKPDLGYFFDDHVRFMDAFIEALGLEEVVLVIHDWGSALGF
HWAKRNPERVKGIAFMEFIRPIPTWDEWPEFARETFQAFRTTDVGRKLIIDQNVFIEGT
LPMGVVRPLTEVEMDHYREPFLNPVDREPLWRFPNELPIAGEPANIVALVEEYMDWL
HQSPVPKLLFWGTPGVLIPPAEAARLAKSLPNCKAVDIGPGLNLLQEDNPDLIGSEIAR
WLSTLEISGGSGSGSKDNTVPLKLIALLANGEFHSGEQLGETLGMSRAAINKHIQTLR
DWGVDVFTVPGKGYSLPEPIPLLNAKQILGQLDGGSVAVLPVVDSTNQYLLDRIGELK
SGDACIAEYQQAGRGSRGRKWFSPFGANLYLSMFWRLKRGPAAIGLGPVIGIVMAEA
LRKLGADKVRVKWPNDLYLQDRKLAGILVELAGITGDAAQIVIGAGINVAMRRVEES
VVNQGWITLQEAGINLDRNTLAATLIRELRAALELFEQEGLAPYLPRWEKLDNFINRP
VKLIIGDKEIFGISRGIDKQGALLLEQDGVIKPWMGGEISLRSAEKGSGSGSHHHHHH
His-tag, HaloTag, TurboID, GS linker

HaloTag-BioID:
MGSEIGTGFPFDPHYVEVLGERMHYVDVGPRDGTPVLFLHGNPTSSYVWRNIIPHVA
PTHRCIAPDLIGMGKSDKPDLGYFFDDHVRFMDAFIEALGLEEVVLVIHDWGSALGF
HWAKRNPERVKGIAFMEFIRPIPTWDEWPEFARETFQAFRTTDVGRKLIIDQNVFIEGT
LPMGVVRPLTEVEMDHYREPFLNPVDREPLWRFPNELPIAGEPANIVALVEEYMDWL
HQSPVPKLLFWGTPGVLIPPAEAARLAKSLPNCKAVDIGPGLNLLQEDNPDLIGSEIAR
WLSTLEISGGSGSGSKDNTVPLKLIALLANGEFHSGEQLGETLGMSRAAINKHIQTLR
DWGVDVFTVPGKGYSLPEPIQLLNAKQILGQLDGGSVAVLPVIDSTNQYLLDRIGELK
SGDACIAEYQQAGRGGRGRKWFSPFGANLYLSMFWRLEQGPAAIGLSLVIGIVMAEV
LRKLGADKVRVKWPNDLYLQDRKLAGILVELTGKTGDAAQIVIGAGINMAMRRVEE
SVVNQGWITLQEAGINLDRNTLAAMLIRELRAALELFEQEGLAPYLSRWEKLDNFIN
RPVKLIIGDKEIFGISRGIDKQGALLLEQDGIIKPWMGGEISLRSAEKGSGSGSHHHHH
H
His-tag, HaloTag, BioID, GS linker

TurboID:
MKDNTVPLKLIALLANGEFHSGEQLGETLGMSRAAINKHIQTLRDWGVDVFTVPGK
GYSLPEPIPLLNAKQILGQLDGGSVAVLPVVDSTNQYLLDRIGELKSGDACIAEYQQA
GRGSRGRKWFSPFGANLYLSMFWRLKRGPAAIGLGPVIGIVMAEALRKLGADKVRV
KWPNDLYLQDRKLAGILVELAGITGDAAQIVIGAGINVAMRRVEESVVNQGWITLQE
AGINLDRNTLAATLIRELRAALELFEQEGLAPYLPRWEKLDNFINRPVKLIIGDKEIFGI
SRGIDKQGALLLEQDGVIKPWMGGEISLRSAEKGSGSGSHHHHHH
His-tag, TurboID, GS linker

BioID:
MKDNTVPLKLIALLANGEFHSGEQLGETLGMSRAAINKHIQTLRDWGVDVFTVPGK
GYSLPEPIQLLNAKQILGQLDGGSVAVLPVIDSTNQYLLDRIGELKSGDACIAEYQQAG
RGGRGRKWFSPFGANLYLSMFWRLEQGPAAIGLSLVIGIVMAEVLRKLGADKVRVK
WPNDLYLQDRKLAGILVELTGKTGDAAQIVIGAGINMAMRRVEESVVNQGWITLQEA
GINLDRNTLAAMLIRELRAALELFEQEGLAPYLSRWEKLDNFINRPVKLIIGDKEIFGIS
RGIDKQGALLLEQDGIIKPWMGGEISLRSAEKGSGSGSHHHHHH
His-tag, BioID, GS linker

### Materials and Methods

#### Construction of plasmids

For cloning, PCR fragments of HaloTag-APEX2, HaloTag-BioID, HaloTag-TurboID, BioID and TurboID with a 6 × His tag at the C-terminus were amplified using Phanta polymerase (Vazyme, P520-01). Then, the fragments were cloned into pET21a vectors by homemade Gibson assembly reagents.

#### Expression of recombinant proteins

*E. coli* BL21 (DE3) cells were transformed with HaloTag-APEX2, HaloTag-BioID, HaloTag-TurboID, BioID and TurboID constructs. Then these cells were cultured in 500 mL of LB medium containing 100 μg/mL of ampicillin at 37°C for 4 h. Once the value of OD600 reached 0.5, as measured by a Tecan plate reader, protein expression was induced by the addition of 0.5 mM IPTG. Subsequently the cultures were incubated at 16°C for 20 h. The cells were harvested by centrifugation at 5,000 rpm for 10 min. After resuspending in binding buffer (50 mM Tris, 300 mM NaCl, pH 7.5), the cells were lysed by sonication for 30 min. The lysate was then centrifuged at 11,000 g for 30 min at 4°C to collect the supernatant, which was then incubated with a gravity column containing 2 mL of Ni-NTA agarose beads. Excess liquid was removed by gravity flow. The beads were then washed sequentially with wash buffers, first with a buffer containing 10 mM imidazole, followed by one with 20 mM imidazole. The recombinant protein was eluted using an elution buffer (50 mM Tris, 300 mM NaCl, pH 7.5, 200 mM imidazole). The purified protein was concentrated using a 10 kDa ultrafiltration tube, and the buffer was exchanged for phosphate-buffered saline (PBS). Finally, the proteins were aliquoted and stored at -80°C for further use.

#### Preparation of staple strands modified with phenol group

The amino-modified ssDNA was ordered from Sangon Biotech, and dissolved in 0.1 M $Na_2B_4O_7$ (pH 8.5) to a concentration of 100 μM. It was then mixed with 100 equivalents of NHS-phenol (CAS: 34071-95-9, dissolved in DMSO). The mixture was shaken at 1000 rpm overnight at room temperature. The phenol-modified ssDNA was recovered using an ethanol precipitation method. Specifically, 0.1 volume of 3 M sodium acetate (pH 5.2) was added to the reaction solution, followed by 2.5 volumes of anhydrous ethanol. The mixture was placed on ice for 30 min and subsequently centrifuged at 12,000 rpm for 30 min at 4°C. The supernatant was carefully discarded, and the pellet was washed twice with 70% ethanol. The tube cap was left open for 5 min to allow residual alcohol to evaporate. The pellet was resuspended in ultrapure water. Finally, the phenol-modified ssDNA was verified by mass spectrometry (LTQ XL™ Linear Ion Trap Mass Spectrometer, Thermo) and stored at -20°C before use.

#### Preparation and purification of DNA origami

The rectangular DNA origami sequence was based on the research in 2006[1]. The staple strands were purchased from Sangon Biotech. M13mp18 scaffold strand was produced in-house according to previous work[2,3]. A 50 μL solution was prepared, containing M13mp18 scaffold strand (10 nM), phenol-modified staple strands (200 nM), APEX2 anchoring strand (200 nM), internal reference strand (200 nM) and other staple strands (100 nM), all in a TAE-$Mg^{2+}$ buffer (40 mM Tirs-acetic acid, pH 8.0, 2 mM EDTA, 12.5 mM $MgCl_2$). The solution was annealed using the following program: 95°C for 15 min, then cooled to 20°C at a rate of 1°C/min. The assembled origami was purified with PEG-8000 to remove the excess staple strand[4]. Specifically, an equal volume (50 µL) of precipitation buffer (15% PEG-8000 (w/v), 5 mM Tris, pH 8.0, 1 mM EDTA, and 505 mM NaCl) was added. The mixture was then placed at 4°C for 10 min, and centrifuged at 5000 g for 20 min. The supernatant was gently discarded and the pellet was resuspended in 50 µL of TAE-$Mg^{2+}$ buffer. The above process was repeated twice, the pellet was resuspended in 50 µL of TAE-$Mg^{2+}$ buffer and stored at -20°C.

**Preparation of APEX2-ssDNA**

First, we prepared ssDNA conjugated with a HaloTag ligand. NHS-Alkynyl (CAS: 1858242-47-3) and $NH_2$-HaloTag ligand (CAS: 1035373-85-3) were mixed in DMSO at a ratio of 1:10 and reacted at room temperature for 30 min. Then, ssDNA-azide (Poly-LGT-$N_3$) was added to the mixture. A solution (50 µL) containing ssDNA-azide (100 µM), Alkynyl-NHS (1 mM), $NH_2$-HaloTag ligand (10 mM), THPTA (2 mM), $CuSO_4$ (1 mM), and sodium ascorbate (5 mM) was reacted at room temperature for 1 h at 1000 rpm. Ethanol precipitation was then performed according to the previous method to prepare the ssDNA-HaloTag ligand. The conjugate was resuspended in PBS and stored at -20°C.

Subsequently, ssDNA-HaloTag ligand (100 µM) and APEX2-HaloTag (50 µM) were mixed and reacted at room temperature for 1 h. The mixture was purified using a Ni-NTA column to remove unreacted ssDNA-HaloTag ligand, followed by ion exchange chromatography to separate out any unreacted proteins. For ion exchange chromatography, the APEX2-ssDNA conjugate was concentrated using a 10 kD ultrafiltration tube and the buffer was exchanged with buffer A (50 mM Tris, pH 7.5). The mixture was then loaded onto an anion exchange column in ÄKTA pure™ chromatography system and eluted using gradient of buffer B (50 mM Tris, pH 7.5, 1 M NaCl), with the gradient increasing from 0–30% over 5 min, then 40–70% over 30 min, and finally 70–100% over 5 min. The APEX2-ssDNA conjugate product was concentrated and buffer-exchanged to PBS using a 10 kD ultrafiltration tube. The conjugate was stored at -80°C before use.

**Assembly and purification of APEX2-origami**

The APEX2-ssDNA conjugate was incubated with DNA origami at room temperature for 30 min, with the conjugate at a fivefold excess. Unbound APEX2-ssDNA was removed using a 100 kD ultrafiltration tube by centrifuging it 5 times at 4000 g for 10 min each. Finally, the buffer was exchanged with a reaction buffer (PBS, 12.5 mM $MgCl_2$) using an ÄKTA pure™ chromatography system, and the sample was stored at -20°C before performing subsequent reactions.

**AFM imaging**

For topographic AFM imaging under liquid conditions, a 0.5 nM sample of 15 µl was deposited onto a freshly cleaved mica surface for 1 min followed by adding 100 mM $NiCl_2$ of 1.5 µl onto the mica for 30 s. Silicon nitride probe (ScanAsyst-Fluid+, Bruker) with a tip radius of 2 nm and a nominal spring constant of 0.7 N/m was typically used. Imaging was performed at room temperature in peak force tapping mode on BioScope Resolve AFM (Bruker) with a resolution of 256 x 256 pixels at a scan rate of 0.5 Hz. A Peak Force amplitude of 25 nm was used, and the force imparted on the sample during imaging was 80 pN. All the AFM images were processed and analyzed with NanoScope Analysis software (Bruker, version 1.8).

**Verification of qPCR specificity**

To verify the specificity of primer A for a specific dsDNA A, all the functional strands were mixed and recorded as sample 1. dsDNA A was removed from sample 1, creating sample 2. Both sample 1 and sample 2 were subjected to qPCR using P141 and specific primer A to observe the amplification and melting curves.

#### Workflow:

**Step 1: Proximity labelingusing APEX2-DNA origami system**

To initiate the reaction, 500 μM biotin-phenol and 1 mM hydrogen peroxide were added to the prepared APEX2-origami sample and allowed to react for 30 min at room temperature. The reaction was terminated by the addition of 10 mM trolox and 10 mM sodium ascorbate. The mixture was then purified using PEG-8000 precipitation[4]. The pellet was resuspended in 20 μL of TAE-$Mg^{2+}$ buffer (40 mM Tris-acetic acid, pH 8.0, 2 mM EDTA, 12.5 mM $MgCl_2$).

**Step 2: Displacement and purification**

The staple strand was separated from the DNA origami by strand displacement. To the resuspended sample in step 1, 2 μL of primer F-5.2, 1 μL of Deep Vent DNA polymerase (NEB, M0259), and 5 μL of 10× buffer were added, with water added to reach a total volume of 50 μL. The reaction proceeded at 72°C for 1 h. The reaction sample was purified and separated using the DNA cleanup kit (NEB, T1030). Briefly, the mixture (50 μL) was combined with 200 μL of Binding buffer, loaded to a column (in DNA cleanup kit), and centrifuged for 1 min at 13,000 rpm to collect the filtrate. Then, 50 μL of water and 600 μL of anhydrous ethanol were added to the filtrate. The mixture was transferred to a new column and centrifuged for 1 min at 13,000 rpm. The column was washed twice with Wash buffer and finally eluted with 20 μL of ultrapure water.

**Step 3: DNA extension**

We first optimize the condition of staple extension. The functional staple strands were mixed, and 2 μL of TL141-AA (10 μM) and 40 μL of 2× Phanta Max (Vazyme, P520-01) were added to the mixture. The mixture was then extended for 1 min at each of the following temperatures: 35°C, 40°C, 50°C, 60°C, and 72°C. The sample was then purified using a DNA cleanup kit and verified by a 10% PAGE gel.

To the purified sample in step 2, 2 μL of TL141-AA (10 μM) and 40 μL of 2× Phanta Max were added and then extended at 50°C for 1 min. The mixture was purified using DNA cleanup kit and finally eluted with 20 μL of ultrapure water, recorded as Input.

**Step 4: DNA enrichment**

Streptavidin beads were blocked with a blocking buffer (1 μg/μL random DNA and 1 μg/μL BSA) for 2 h. Then, 0.6 μL of blocked streptavidin beads and 14.4 μL of 2× loading buffer (100 mM Tris, pH 7.5, 1 M NaCl, 10 mM EDTA, 0.2% v/v Tween-20) were added to 15 μL of the Input sample and then the mixture was incubated at 1000 rpm for 45 min. After removing the supernatant, the beads were washed three times with washing buffer (100 mM Tris, pH 7.5, 4 M sodium chloride, 10 mM EDTA, 0.2% v/v Tween-20) at room temperature, twice with washing buffer at 50°C for 10 min, and twice with PBS. The beads were resuspended in 20 μL of elution buffer (95% v/v formamide, 10 mM EDTA, pH 8.0, 1.5 mM biotin) and incubated at 95°C for 10 min. The supernatant was purified using DNA cleanup kit and recorded as Enrich.

**Step 5: qPCR**

Both the Input and Enrich samples were subjected to qPCR (instrument: CFX96, Bio-Rad) using primer P141 and specific primers with the Takara qPCR kit (RR820A). The Ct values of each staple and internal reference were recorded to calculate the relative labeling efficiency of each site using **Equation 1**.

**Determination and calculation of amplification efficiency of each staple strand**

The Input sample was diluted with EASY Dilution (Takara, 9160), then the diluted samples underwent qPCR with P141 and corresponding primers. The fitting curve of N and Ct values was drawn according to the formula: $Lg(N) = -Ct * lg(1+E) + C$, and then the amplification efficiency E was calculated.

**Preparation and purification of scaffold strands-HaloTag-APEX2, scaffold strands-HaloTag-TurboID, and scaffold strands-HaloTag-BioID**

Scaffold strands modified with 3'-amino and 5'-amino groups was ordered from Sangon Biotech, and dissolved in 0.1 M $Na_2B_4O_7$ (pH 8.5) and mixed in equal molar amounts. To the mixture, 100 equivalents of NHS-Alkynyl (CAS: 1858242-47-3, dissolved in DMSO) were added and reacted at room temperature overnight at 1000 rpm. The modified scaffold strands were purified by ethanol precipitation as previously described, with the pellet resuspended in PBS and stored at -20°C until needed.

HaloTag ligand conjugation with scaffold strands was performed via a click reaction. A solution (50 μL) containing alkynyl-modified scaffold strands (100 μM), azide-HaloTag ligand (CAS: 2568146-55-2, 10 mM), THPTA (2 mM), $CuSO_4$ (1 mM), and sodium ascorbate (5 mM) was reacted at room temperature for 1 h at 1000 rpm. The mixture was purified by ethanol precipitation as previously described. The conjugate was resuspended in PBS and stored at -20°C.

Subsequently, scaffold strands conjugated with the HaloTag ligand (100 μM) were mixed with APEX2-HaloTag (50 μM), TurboID-HaloTag (50 μM) or BioID-HaloTag (50 μM) and reacted at room temperature for 1 h. Unreacted ssDNA-HaloTag ligand was removed using a Ni-NTA column, followed by further purification of the sample via ion exchange chromatography as previously described. The conjugates were stored at -80°C until needed.

**Measurement of TurboID and BioID labeling in ds-DNA system**

The amino-modified staple strands were mixed with three equivalents of either scaffold strands-HaloTag-TurboID or scaffold strands-HaloTag-BioID. The mixture was incubated at room temperature for 30 min, after which the buffer was replaced with 0.1 M $Na_2B_4O_7$ (pH 8.5) + 12.5 mM $Mg^{2+}$ using a desalting column in ÄKTA pure™ chromatography system. The final concentration of each amino-modified staple strand was 1 nM, and the proximity labeling was initiated by adding 1 mM ATP and 1 mM Biotin. The reaction was allowed to proceed for 30 min, 2 h, and 12 h. The reaction was quenched with 10 mM Tris-HCl, followed by the addition of 300 μL of formamide to denature the dsDNA into ssDNA. Subsequently, the mixture was purified using a DNA cleanup kit and eluted with 20 μL of ultrapure water.

Next, 2 μL of TL141-AA (10 μM) and 40 μL of 2 × Phanta max were added to the purified sample and extended at 50°C for 1 min. Then, 200 μL of lysis buffer (PBS, 300mM NaCl) was added, and then scaffold strands-HaloTag-TurboID or scaffold strands-HaloTag-BioID were removed using a Ni-NTA column, with the filtrate collected. The filtrate was purified with a DNA cleanup kit and eluted with 20 μL of ultrapure water, recorded as Input.

The pull-down and qPCR steps are the same as those used in the APEX2-origami system.

**Synthesis of Biotin-5'-AMP**

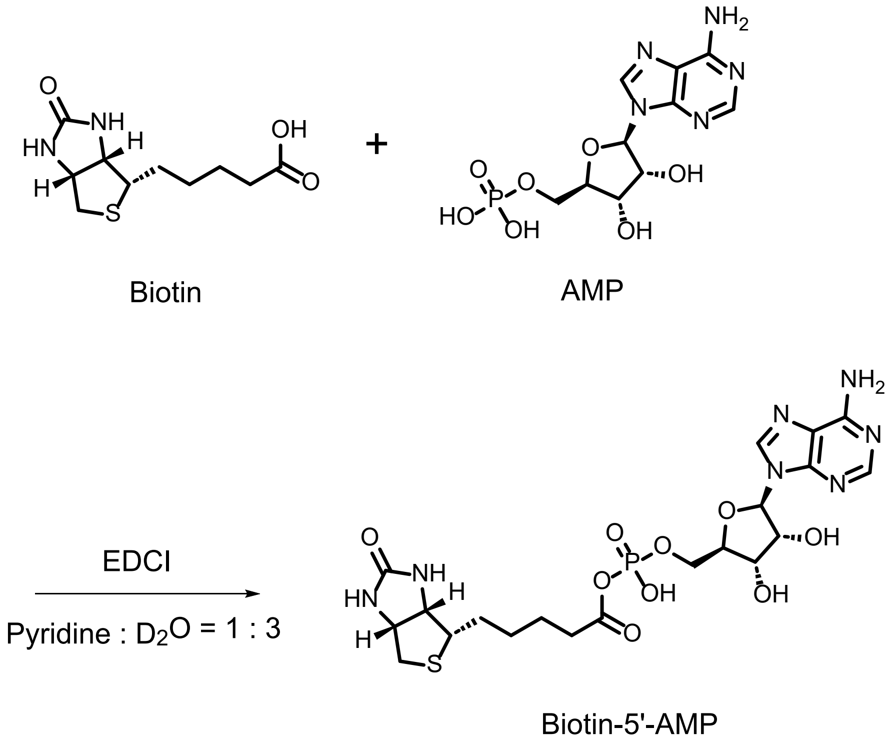


Biotin (24.4 mg, 0.10 mmol, 1.0 eq) and adenosine monophosphate (AMP, 38.2 mg, 0.11 mmol, 1.1 eq) were dissolved in 0.4 mL of a solution containing pyridine and $D_2O$ in a 1:3 ratio at 0°C. 1-(3-Dimethylaminopropyl)-3-ethylcarbodiimide hydrochloride (EDCI, 76.7 mg, 0.40 mmol, 4.0 eq) was then added to the mixture. The mixture was stirred at 0°C for 3 hours until the reaction was complete. The progress of the reaction was monitored by LC/MS (4.6 mm × 150 mm 3 μm C18 column; 2 μL injection; 5-100% $CH_3CN/H_2O$, linear gradient, with constant 0.1% v/v formic acid additive; 6.5 min run; 0.4 mL/min flow; ESI; positive ion mode; UV detection with

ACQUITY PDA). The reaction mixture was then purified by HPLC (eluent, a 45 min linear gradient, from 10% to 100% solvent B; flow rate, 10.0 mL/min; detection wavelength, 254 nm; eluent A (ddH2O containing 0.1% trifluoroacetic acid (v/v)) and eluent B ($CH_3CN$). Column: Orienda Prep C18 column, FLA-2015-5-120A-BP, 120 Å, 5 µm, 20 x 150 mm). **Biotin-5'-AMP** (0.14 mmol, 69.8% yield) was finally obtained as a white solid upon freeze-drying.
ESI-MS: m/z calculated for $C_{20}H_{29}N_7O_9PS^+$, **Biotin-5'-AMP** $(M+H)^+$ 574.2, found 574.4.
LC trace:

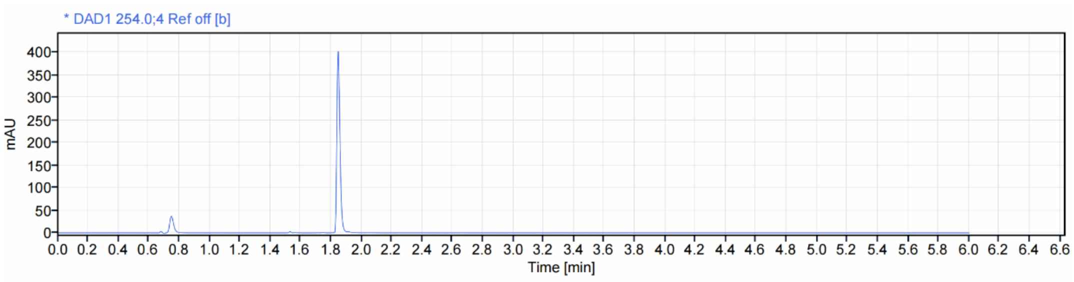


**Determination of dissociation constant and binding constant by stopped-flow spectroscopy**

The rates of association and dissociation were measured by stopped-flow spectroscopy using a SX20 stopped-flow instrument (Applied Photophysics) following the methods of previous work[5]. In detail, the excitation wavelength was set at 280 nm for all measurements, and fluorescence emission was monitored above 340 nm using a cutoff filter. For association of biotin-5'-AMP and TurboID or BioID, both TurboID or BioID in Standard Buffer (10 mM Tris-HCl, pH 7.5, 200 mM KCl, 2.5 mM $MgCl_2$) and a solution containing biotin-5'-AMP were rapidly mixed at a 1:1 ratio. Subsequently, the time-dependent change in the intrinsic protein fluorescence was measured. The final concentration of TurboID or BioID was 1 µM and the concentrations of biotin-5'-AMP were varied ranged from 5 µM to 15 µM to satisfy pseudo-first order conditions. Stopped-flow fluorescence traces obtained are well described by the following double exponential equation S1, where F is the measured fluorescence intensity, and *A1* and *τ1* and *A2* and *τ2* are the amplitudes and relaxation times for the first and second kinetic phases, respectively. We obteined *τ1* and *τ2* at different concentrations of biotin-5'-AMP according to **Equation S1**. We then calculated the dissociation constant $k_1$ using **Equation S2**.

$$F = A_1 \exp(t \times 1/\tau_1) + A_2 \exp(t \times 1/\tau_2) + C$$

**Equation S1**

$$\frac{1}{\tau_1} + \frac{1}{\tau_2} = k_1[\text{ligand}] + k_{-1} + k_2 + k_{-2}$$

**Equation S2**

For dissociation of biotin-5'-AMP and TurboID or BioID, preformed complexes of TurboID·biotin-5'-AMP or BioID·biotin-5'-AMP in Standard Buffer were rapidly mixed with a solution containing 100 µM biotin in a 1:1 ratio. The subsequent time-dependent decrease in fluorescence was then measured. The resulting kinetic traces were analyzed using a single exponential model (**Equation S3**) and **Equation S4** to obtain unimolecular dissociation constants $k_{off}$.

$$F = A \exp\left(\frac{t}{\tau}\right) + C.$$

**Equation S3**

$k_{off} = 1 / \tau$ **Equation S4**

Finally, we calculate $K_D$ using **Equation S5**.

$K_D = k_{off} / k_1$ **Equation S5**
